# Extracting a low-dimensional description of multiple gene expression datasets reveals a potential driver for tumor-associated stroma in ovarian cancer

**DOI:** 10.1101/048215

**Authors:** Safiye Celik, Benjamin A Logsdon, Stephanie Battle, Charles W Drescher, Mara Rendi, Hawkins David R, Su-In Lee

## Abstract

**Background:** Discovering patient subtypes and molecular drivers of a subtype are difficult and driving problems underlying most modern disease expression studies collected across patient populations. Expression patterns conserved across multiple expression datasets from independent disease studies are likely to represent important molecular events underlying the disease.

**Methods:** We present the INSPIRE (<u>**IN**</u>ferring <u>**S**</u>hared modules from multi<u>**P**</u>le gene exp<u>**RE**</u>ssion datasets) method to infer highly coherent and robust *modules* of co-expressed genes and the dependencies among the modules from multiple expression datasets. Focusing on inferring modules and their dependencies conserved across multiple expression datasets is important for several reasons. First, using multiple datasets will increase the power to detect robust and relevant patterns (modules and dependencies among modules). Second, INSPIRE enables the use of multiple datasets that contain different sets of genes due to, e.g., the difference in microarray platforms. Many methods designed for expression data analysis cannot integrate multiple datasets with variable discrepancy to infer a single combined model, whereas INSPIRE can naturally model the dependencies among the modules even when a large proportion of genes are not observed on a certain platform.

**Results:** We evaluated INSPIRE on synthetically generated datasets with known underlying network structure among modules, and gene expression datasets from multiple ovarian cancer studies. We show that the model learned by INSPIRE can explain unseen data better and can reveal prior knowledge on gene functions more accurately than alternative methods. We demonstrate that applying INSPIRE to nine ovarian cancer datasets leads to the identification of a new marker and potential molecular driver of tumor-associated stroma - *HOPX*. We also demonstrate that the *HOPX*module strongly overlaps with the genes defining the mesenchymal patient subtype identified in The Cancer Genome Atlas (TCGA) ovarian cancer data. We provide evidence for a previously unknown molecular basis of tumor resectability efficacy involving tumor-associated mesenchymal stem cells represented by *HOPX*.

**Conclusions:** INSPIRE extracts a low-dimensional description from multiple gene expression data, which consists of modules and their dependencies. The discovery of a new tumor-associated stroma marker, *HOPX,* and its module suggests a previously unknown mechanism underlying tumor-associated stroma.

## BACKGROUND

### Introduction

As datasets increase in size, scope, and generality, the possibility to infer potentially *relevant* and *robust* features from data increases. Extracting a biologically intuitive low-dimensional representation (LDR) of data in an *unsupervised* fashion (i.e., based on the underlying structure in the data, not with respect to a particular prediction task) has become an important step to identify robust and relevant information from data. Development of unsupervised LDR learning methods is a very active area of modern research in machine learning and high dimensional data analysis^1–3^. Specific machine learning domains to see noted success recently include the development of deep learning algorithms^3^, where authors demonstrate enormous increases in performance on difficult tasks such as image and text classification^4,5^. Analogously, in cancer transcriptomics unsupervised LDR learning has seen success on very difficult problems, such as predicting patient outcome in breast cancer in the DREAM7 breast cancer prognosis challenge^6^. The winning team leveraged an unsupervised LDR extraction method on independent transcriptomic data from multiple cancer types, and significantly outperformed the other contestants in the challenge by a large margin^7^ along with all other known prognostic signatures in breast cancer.

There are three main challenges with applying existing unsupervised LDR learning approaches to cancer transcriptomic data. First, any one study may not be generalizable in that there will be either technical (e.g. sample ascertainment) or experimental (e.g. batch effects) confounders that make an LDR of data extracted from an individual dataset in a naïve way not necessarily generalizable to other datasets. Second, identifying simple modules (co-expressed sets of genes) using methods such as WGCNA^8^ or simple clustering approaches^9,10^ will not necessarily capture complex dependence structures among the modules. Appropriately accounting for rich dependencies among these modules will improve their biological coherence. It has been shown that modeling the dependencies among modules improves the quality of the inferred modules from gene expression data^11^. Finally, and most importantly, most cancer transcriptomic data is within the *p* » *n* regime (high-dimensional), i.e. we usually have tens of thousands of genes, but only hundreds of samples at most. This means that a successful method must include a very aggressive dimensionality reduction mechanism that allows generalization across datasets, since the potential for overfitting is high. This implies that models that allow for arbitrarily rich dependencies among variables (such as those used in deep learning methods) cannot necessarily be applied without overfitting the data.

We present a novel unsupervised LDR learning method, called INSPIRE (<u>**IN**</u>ferring <u>**S**</u>hared modules from multi<u>**P**</u>le gene exp<u>**RE**</u>ssion datasets), to infer highly coherent and robust modules of genes and their dependencies on the basis of gene expression datasets from multiple independent studies (Figure 1).INSPIRE is an unconventional and aggressive data dimensionality reduction approach that extracts highly biologically relevant and coherent modules from gene expression data, where the number of samples is much less than the number of observed genes – the norm for cancer expression data. INSPIRE addresses the three aforementioned challenges. First, INSPIRE naturally integrates many datasets by modeling the latent (hidden, unobserved) variables in a probabilistic graphical model^12^, where the latent variables are modeled as a Gaussian graphical model, the most commonly used probabilistic graphical model for continuous-valued variables (Figure 1). Each observed gene is treated as a noisy and independent observation of these underlying latent variables. By jointly inferring the assignment of observed genes to latent variables and the structure of the Gaussian graphical model among these latent variables, we can naturally capture both modules and their dependencies that generalize across multiple datasets (Figure 1). This addresses the issue with generalizability of modules across datasets. Second, our method naturally models the dependencies among the modules, which allows us to capture more complicated dependencies among pathways, cell populations, or other biologically driven modules than naïve approaches such as hierarchical clustering. In a previous study^11^, we have shown that modeling the dependencies among modules directly improves the biological coherence of the modules we learn, and their generalizability across datasets. Finally, by modeling the data as noisy observations from a much lower dimensional subset of modules, we are able to overcome the curse of dimensionality, and have better power to both learn the modules and their dependencies, even when the number of genes is much greater than the samples size. Through extensive simulated and real data analysis (Figure 2) we demonstrate our approach is a great practical tradeoff between model complexity and model parsimony when understanding biological pathways characterizing the cancer transcriptome across ovarian cancer patients.

**Figure 1.**
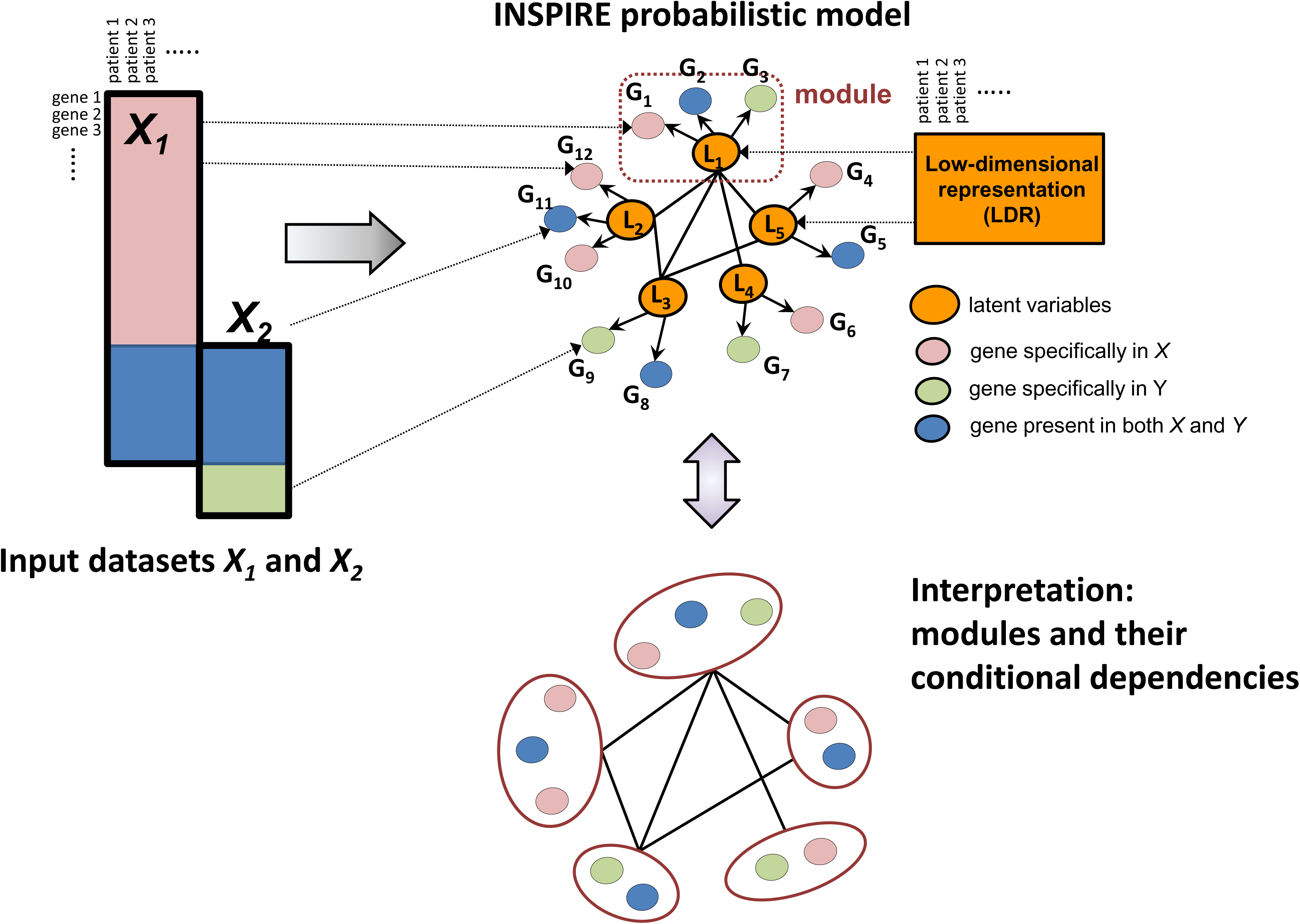
Overview the INSPIRE framework. INSPIRE takes as input multiple expression datasets that potentially contain different sets of genes and learns a network of expression modules (i.e., co-expressed sets of genes) conserved across these datasets. INSPIRE is a general framework that can take any number of datasets as input; two datasets (*X*_*1*_ and *X*_*2*_) are shown in representation for simplicity. **Top left:** Two input datasets are represented by rectangles with black solid lines. Rows represent genes and columns represent samples. The blue region contains the data for the genes that are contained in both datasets. The pink and green regions contain the data for the genes which are contained by only one of the datasets. **Top right:** The features (latent variables), each corresponding to a module, are shown by the orange matrix as learned by INSPIRE. These are used as a low-dimensional representation of the expression datasets. **Top middle:** As an example, five INSPIRE features *L*_1_,…,*L*_5_ (orange-shaded circles), 12 genes *G*_1_,…,*G*_12_ associated with those features, and the conditional dependency network among the INSPIRE features are represented. The dependencies among features are conserved across the datasets. **Bottom middle:** Five modules, each corresponding to an INSPIRE feature, and the dependency network among them are represented as the interpretation of the INSPIRE features and their conditional dependencies.

**Figure 2.**
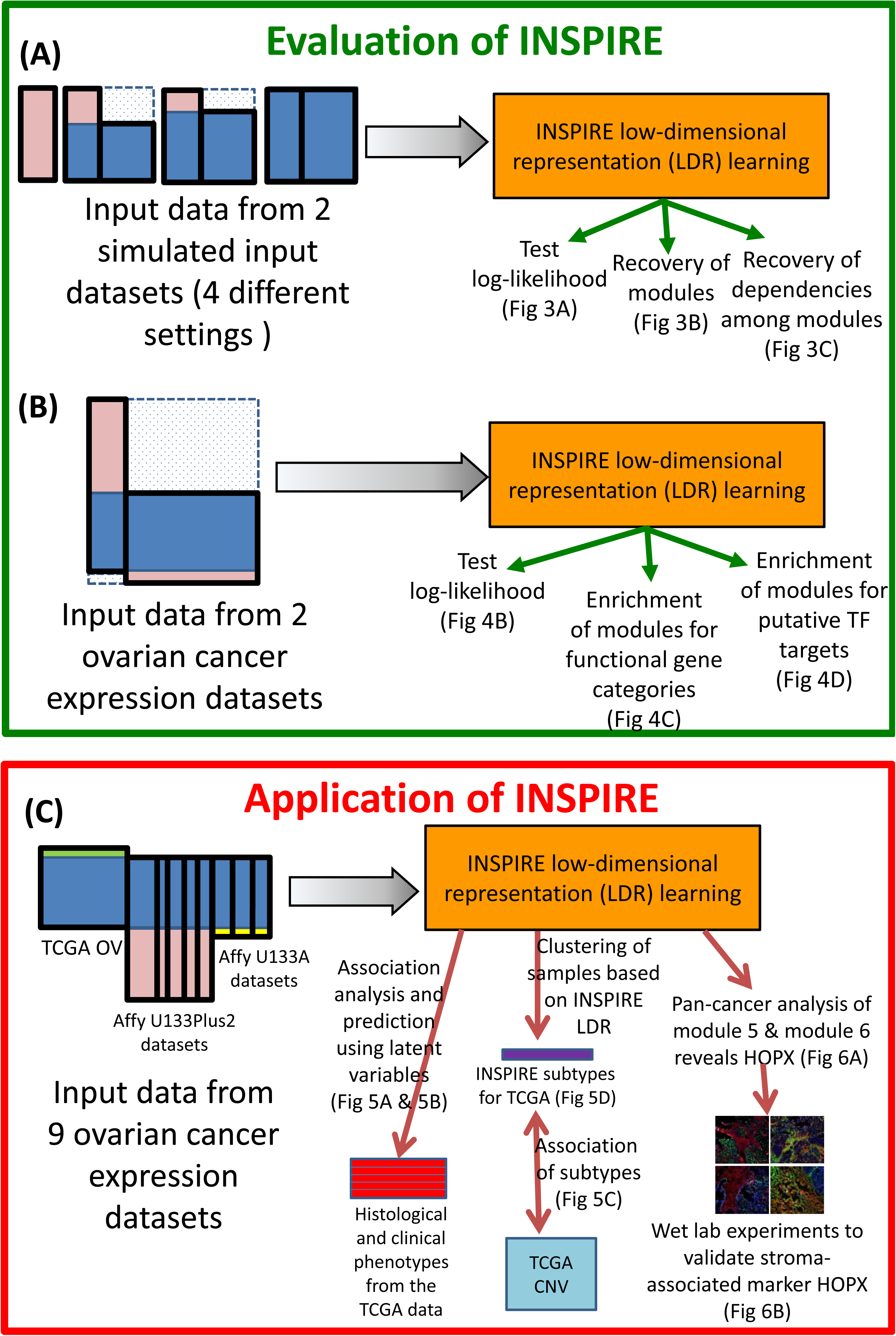
Overview of the evaluation and application of INSPIRE procedure. The procedure takes as input *K* ≥ 2datasets, and the method is an iterative procedure that determines both the assignment of the genes to modules, the features each corresponding to a module, and the dependencies among the features which are conserved across the datasets. **(A)** Evaluation of INSPIRE using simulated data. Two simulated datasets in four settings corresponding to different amount of gene overlap are provided as input to the INSPIRE learning algorithm, and the learned modules and network are evaluated in terms of three different metrics. **(B)** Evaluation of INSPIRE using two ovarian cancer expression datasets. Two expression datasets from different platforms are provided as input to the learning algorithm and the learned modules and network are evaluated in terms of three different metrics. **(C)** Application of INSPIRE on nine real-world ovarian cancer expression datasets. As an application of INSPIRE, we first check the association of the learned INSPIRE features with six histological and clinical phenotypes, which is followed by subtyping the patients into groups based on the learned INSPIRE features. Observing that INSPIRE features have high association with the histological and clinical phenotypes in cancer, and the subtypes learned based on the features can predict CNV abnormalities well leads us to do a deeper analysis of two modules (modules 5 and 6), which are good predictors of many phenotypes and good differentiators of learned ovarian cancer subtypes.

When we apply INSPIRE to nine gene expression datasets from ovarian cancer studies (Figure 2C), we identify a novel tumor-associated stromal marker *HOPX*, which additional analyses suggest may be a molecular driver for a conserved module in the network that contains known epithelial-mesenchymal transition (EMT) inducers and is significantly associated with percent stroma in ovarian tumors from The Cancer Genome Atlas (TCGA). This module is one of the two modules that best represent one of the predominant subtypes of ovarian cancer, ‘mesenchymal’ subtype identified in the TCGA ovarian cancer study^13^. These multiple lines of evidence suggest that *HOPX* may be a great target for further functional validation to understand the maintenance of tumor-associated stroma along with understanding the clinically relevant ‘mesenchymal’ subtype in ovarian cancer.

The implementation of INSPIRE, the data used in the study, and the resulting INSPIRE models are freely available on our website^14^.

### Literature overview

INSPIRE extracts a low-dimensional representation (LDR) of multiple expression datasets, by fitting a probabilistic model with latent variables and their dependencies into the input data (Figure 1). Previous approaches can be divided into two categories; 1) *supervised* methods that extract an LDR that is discriminative of different class labels in the training samples, and 2) *unsupervised* methods (including INSPIRE) that extract an LDR purely based on the underlying structure of the data.

A supervised method aims to extract an LDR that is discriminative between class labels in a particular prediction problem. Several authors developed methods that use known pathways or biological networks along with gene expression data to extract an LDR (“pathway markers”) whose activity is predictive of a given phenotype^15–18^. Chuang et al.^15^ proposed a greedy search algorithm to detect subnetworks in a given protein-protein interaction (PPI) network, such that each subnetwork contains genes whose average expression level is highly correlated with class labels (metastatic/non-metastatic) measured by the mutual information. The authors claim that subnetwork markers outperform individual genes for predicting breast cancer metastasis. Lee et al.^16^ developed a similar algorithm to select subsets of genes from MSigDB (Molecular Signatures Database) C2 (curated) pathways that give the optimal discriminative power for the classification of leukemia/ breast cancer phenotypes. Both Chuang et al.^15^ and Lee et al.^16^ determined LDR as the average expression levels of genes in each subnetwork and pathway, respectively. Taylor et al.^17^ proposed a similar approach that uses a PPI network, but instead of computing the LDR by averaging gene expression levels within a subnetwork (or a pathway), they compute the expression difference between a hub protein and all of its neighbors in the PPI network. Ravasi et al.^18^ used a similar approach to extract subnetwork features as hub TFs from TF PPI networks in human and mouse. Besides the methods that infer an LDR by averaging (or aggregating) expression levels of subsets of genes, there have been methods to select a subset of genes. For example, Herschkowitz et al.^19^ used 106 genes selected by the *intrinsic analysis* for a classification problem (122 mouse breast tumors/232 human breast tumors). The intrinsic analysis aims to select genes that are relevant to tumor classification by identifying genes whose expression show relatively low within-group variation and high between-group variation for known groups of tumors in each of human and mouse datasets^19^. Although supervised methods would be useful to infer an LDR relevant to a particular prediction problem, there are several disadvantages over unsupervised methods. First, we need to have a particular prediction problem with class labels, which may not be available. Second, they usually rely on the assumption that the same genes are differentially expressed in all samples within a class, which is unlikely to be true in heterogeneous diseases such as cancer.

On the other hand, unsupervised LDR learning methods extract an LDR without knowing about the class labels, while the learned LDR can be used for classification purposes later. One of the most commonly used methods is the principal component analysis (PCA)^20^ which sequentially extracts most of the variance (variability) of the data. However, each PC (principal component - or eigengene) is a linear combination of all genes not a small subset of genes, which makes it difficult to biologically characterize it. Clustering algorithms^21^, on the other hand, generate explicit gene clusters, and they define an LDR as a set of mean or median expression levels of the genes in each cluster. In the seminal work by Langfelder and Horvath (a technique called WGCNA)^8^, the adjacencies retrieved from Pearson’s correlation of the expression levels of the gene pairs is transformed into topological overlap measure (TOM), namely network interconnectivity that takes into account the shared neighbors of each gene pair, which is then used in a hierarchical clustering to define modules. While WGCNA^8^ defines its similarity measure (i.e., TOM) based on the *marginal correlations* between genes, other authors have used *partial correlations (conditional dependencies)* to model gene relationships ^11,22,23^. Chandrasekaran et al.^22^ incorporated latent variables into a Gaussian graphical model among individual genes, while Celik et al.^11^ divided variables into modules and learned a module-level dependencies (module graphical lasso - MGL). He et al.^23^ defined an LDR as a set of latent factors, and modeled each latent factor as a linear combination of genes (structured latent factor analysis - SLFA). While similar to Celik et al.^11^ in modeling a higher-level dependency structure, He et al.^23^ does not form explicit clusters. Finally, Cheng et al.^7^ identified 12 metagenes, each of which is a weighted average of the genes that are co-expressed across multiple cancer types. They showed that the prediction model they derived based on these metagenes is highly predictive of survival in breast cancer within the context of the DREAM7 Challenge, leading to the top scoring model^6^.

There are three major differences between INSPIRE and previous approaches. First, none of the previous methods to learn LDR can accommodate multiple datasets containing different sets of genes (e.g., different microarray platforms), while INSPIRE directly addresses this challenge. One naïve way to run previous methods on datasets that contain different sets of genes with a partial overlap is to treat the values on the genes that are not observed in each dataset as missing data. We could use missing value imputation techniques to fill in missing data and learn a single statistical model from the imputed data. However, most imputation methods perform poorly when a large number of values are missing (Figure 1). We demonstrate that INSPIRE outperforms the imputation-based approaches (methods named ‘Imp--’ in Figures 3 and 4). Second, INSPIRE uses a novel probabilistic model that can describe more complex relationships (i.e., conditional dependencies) than pairwise marginal correlations among genes. We show that INSPIRE outperforms a correlation-based method, WGCNA. Finally, INSPIRE uses a novel learning algorithm to make use of all samples in multiple datasets, which increases the statistical power to detect a statistical robust model (Figure 1). Our extensive experiments show that these key properties of INSPIRE lead to biologically more relevant and statistically more robust features than alternative methods.

**Figure 3.**
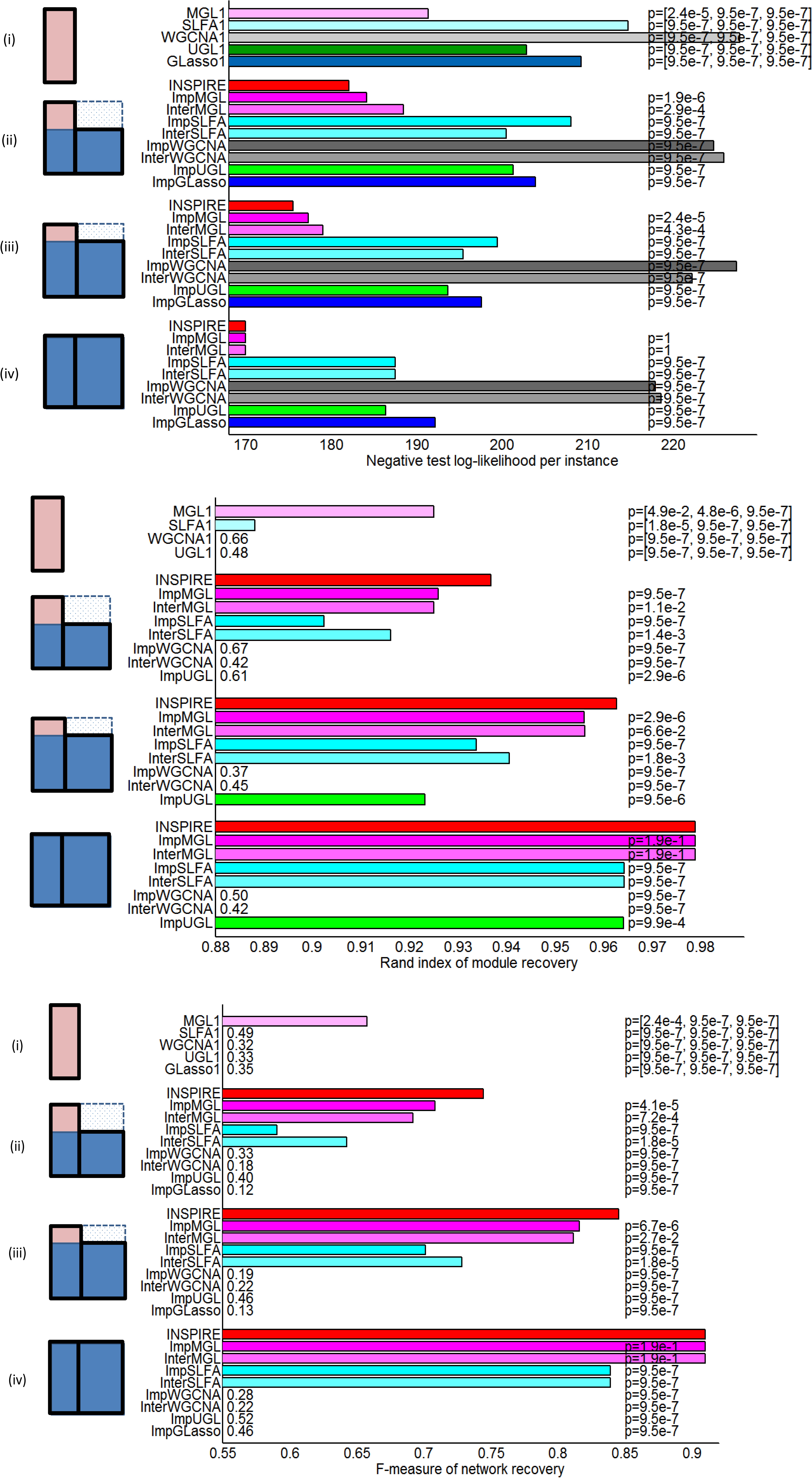
Illustration of the synthetic data, aligned with four groups of bars in each of (A)-(C). Rows represent genes and columns represent samples. **(A)** Negative test log-likelihood per instance averaged over 20 different instantiations of the synthetic data (lower is better). **(B)** Rand index for module recovery averaged over 20 different instantiations of the synthetic data. **(C)** F-measure for feature dependency recovery averaged over 20 different instantiations of the synthetic data. The Wilcoxon signed rank test *p*-value represented on each bar (except the bars for INSPIRE) measures the statistical significance of the difference between the method and INSPIRE.

**Figure 4.**
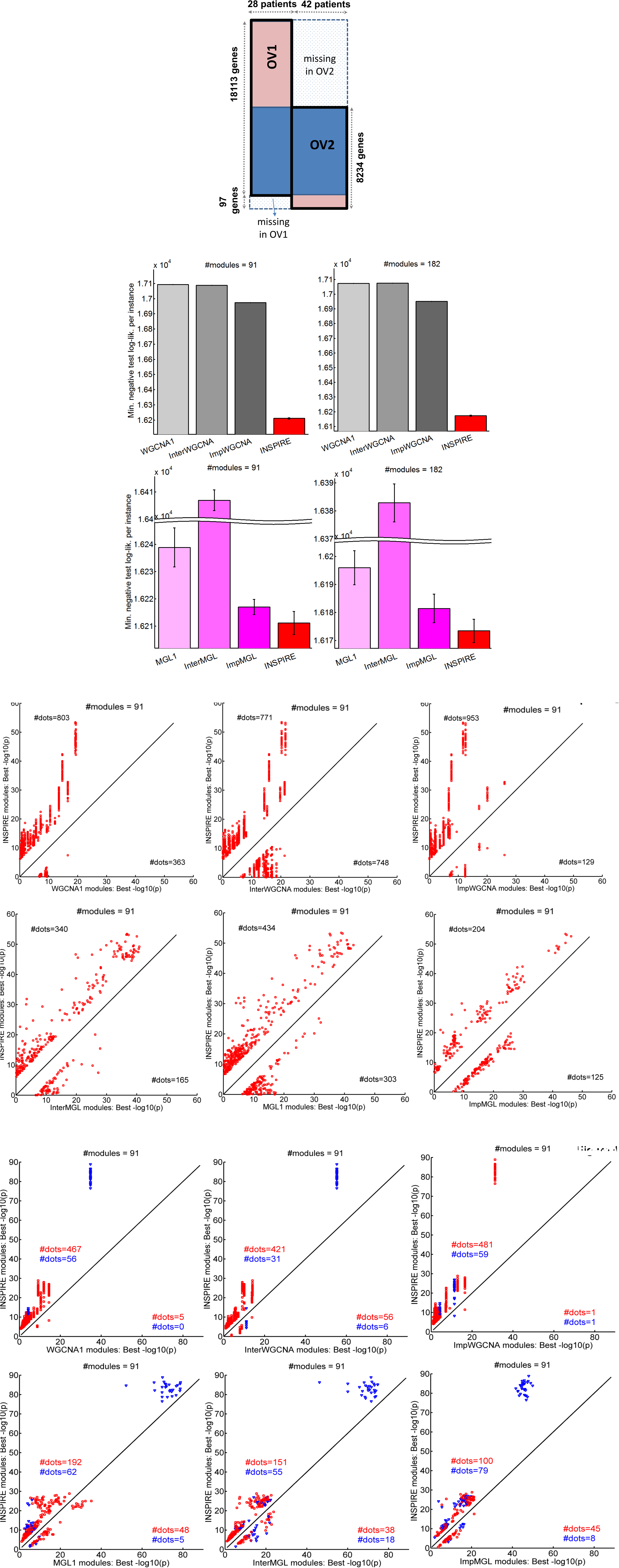
**(A)** Illustration of the two OV datasets used for evaluating INSPIRE. Rows represent genes and columns represent samples. **(B)** For *k* = 91 (left) and *k* = 182 (right), INSPIRE is compared to WGCNA variants (top) and MGL variants (bottom) in terms of the best CV negative test log-likelihood (lower is better) across all tested sparsity tuning parameters (*λ*) **(C)** For *k* = 91, INSPIRE (y-axis) is compared to each of the six competing methods (x-axes) in terms of the best — log_10_ *p* from the functional enrichment of the learned modules. Each dot is a KEGG, Reactome or BioCarta GeneSet, and only the GeneSets with a Bonferroni corrected *p* < .05 in at least one of the compared two methods are shown on each plot. For MGL variants and INSPIRE, results from multiple runs are shown. We only considered the GeneSets with sufficiently different significance between the two methods, i.e., |log_10_ *p*(*INSPIRE*) – log_10_ *p*(*ALTERNATIVE_METHOD*)| ≥ *δ*. *δ* = 6 here and the results were consistent for varying *δ*. **(D)** For *k* = 91, INSPIRE (y-axis) is compared to each of the six competing methods (x-axes) in terms of the best – log_10_ *p* from the ChEA enrichment of the learned modules. Each dot is for a gene set composed of a TF and its targets, and only the sets with a Bonferroni corrected *p* < .05 in at least one of the compared two methods are shown on each plot. For MGL variants and INSPIRE, results from multiple runs are shown. We only considered the TFs with sufficiently different significance between the two methods, i.e., |log_10_ *p*(*INSPIRE*) – log_10_ *p*(*ALTERNATIVE_METHOD*)| ≥ *δ*. *δ* = 3 here and the results were consistent for varying *S*. Each blue dot corresponds to a TF which sits in the INSPIRE module that is significantly enriched for its targets, and each red dot corresponds to a TF which sits in an INSPIRE module different than the one that is significantly enriched for its targets.

## RESULTS

### Overview of the INSPIRE framework

INSPIRE extracts a low-dimensional representation (LDR) from multiple gene expression datasets by inferring *k* latent (unobserved) variables and the dependencies among the latent variables captured by a probabilistic graphical model (Figure 1). INSPIRE uses a standard iterative learning algorithm to optimize the joint log-likelihood objective function, Equation (1), by iteratively updating its model parameters until convergence (see Methods for details). INSPIRE iterates the following three steps until convergence: i) inferring the values of latent variables with all the other parameters held fixed, as described in Equation (3), ii) assigning genes into latent variables as described in Equation (4), and iii) learning a network of latent variables as described in Equation (5). In each iteration, latent variables are computed based on the current assignment of genes into modules and the estimated dependency network among the latent variables, as described in Equation (3). If there are no dependencies among latent variables, each latent variable would be an average expression level of the genes in the module. Thus, latent variables can be viewed as module centers adjusted for the estimated dependency network among latent variables.

A set of genes assigned to the same latent variable is referred to as a *module* (Figure 1). To focus on identifying a parsimonious, independent set of modules from high-dimensional gene expression data, we design our model such that each gene is assigned to only one module, although it would be a simple extension to assign each gene to multiple modules. However, when we implemented an extension of INSPIRE which allows each gene to be assigned to more than one module, the functional coherence of modules significantly decreased (Figure S7). This could be because the model with genes assigned to multiple modules has a significantly increased number of parameters.

The number of modules *k* is determined based on the standard Bayesian Information Criterion (BIC), although users can determine *k* in a different way depending on the problem. INSPIRE framework simultaneously infers the assignment of genes into *k* latent variables and the dependency network among *k* latent variables by fitting the probabilistic model across multiple gene expression datasets that can potentially have different sets of genes (e.g., different platforms) (see Methods). The INSPIRE model provides a biologically intuitive LDR model for gene expression data where many biological networks are *modular* and genes involved in similar functions are likely to be connected more densely with each other. How genes are organized into modules and how these modules are connected with each other would provide improved insights into the underlying disease process, as discussed below.

After evaluating INSPIRE by comparing with alternative methods on simulated data and a small set of genome-wide expression datasets (Figure 2A-B), we applied INSPIRE to many ovarian cancer expression datasets, which lead to a novel marker and potential driver of tumor-associated stroma (Figure 2C).

### INSPIRE learns underlying modules and their dependencies from simulated data more accurately than 13 other methods

We first evaluate INSPIRE on data simulated using a probabilistic model of (unobserved) latent variables, gene expression levels, and the dependencies among the latent variables captured by a probabilistic graphical model (Figure 1). To simulate the situation in which we are given expression datasets that contain different sets of genes (e.g., different microarray platforms), we generated two datasets (Dataset1 and Dataset2) with the same genes and included all genes in Dataset1 and varying percentages of the genes in Dataset2 such that varying numbers of genes are present in the overlapping portion of the datasets. This leads to three settings (Figure S1A, Figure 3A (ii)-(iv) left): (ii) 60% of the genes are present in Dataset2, (iii) 80% of the genes are present in Dataset2; and (iv) all genes are present in Dataset2. The total number of genes in each of these settings is 250, and the number of modules is 10, with an average of 25 genes in a module datasets (see Methods for details of synthetic data generation).

We compare INSPIRE with the following five state-of-the-art methods: **i) <u>GLasso</u>** - standard graphical lasso^24^ that learns a gene-level conditional dependence network with no LDR or module assumption; **ii)** <u>**UGL**</u> - unknown group *L*_1_ regularization^25^ that learn sparse block-structured inverse covariance matrices with unknown block structure; **iii)** <u>**SLFA**</u> - structured latent factor analysis^23^ that learn an LDR of the data as well as the relationship between the latent factors; **iv)** <u>**WGCNA**</u> - weighted gene co-expression network analysis^8^ that allows to define modules based on a special metric derived from the correlations of the gene pairs; **v)** <u>**MGL**</u> - module graphical lasso^11^ which simultaneously learns an LDR and the conditional dependencies among the latent variables (Table 1). Since all those methods work on a single dataset, to enable the application of these methods to multiple datasets with variable discrepancy, we adapt the input data to those five methods in three ways (Figure S1B): 1) using only Dataset1 that contains all genes, 2) using data on the genes that are present in both datasets (blue-shaded region in Figure 1), and assigning the rest of the genes to the learned modules based on the Euclidean distance between the gene’s expression and the expression of each of the modules, and 3) imputing missing values in Dataset2 and using both datasets as if they were a single dataset. This leads to 13 methods (Table 1). InterMGL, ImpMGL and INSPIRE represent different ways of handling missing data: INSPIRE uses a novel learning algorithm that does not require the missing portion when learning; ImpMGL imputes missing variables in the datasets before learning; and InterMGL ignores missing variables in the datasets. We run each method on 20 different instantiations of the synthetic data, and present the average results with *p*-values of significance of the difference with INSPIRE (Methods; Figure 3). We evaluated INSPIRE and 13 competitors in terms of how well they explain unseen data measured by the test-set log-likelihood, gene-module assignment accuracy, and the module dependency network accuracy. In order to make comparison with WGCNA variant methods possible, we applied a standard graphical lasso algorithm to the modules learned by a WGCNA variant method. INSPIRE, SLFA and MGL are iterative algorithms with non-convex objective functions, so their results may depend on the initialization of the parameters. To rule out the possibility to make a conclusion based on a particular set of initial parameters, we performed the variants of those algorithms multiple times with different starting points (see Methods for details on initialization).

**Table 1.** Methods we compared with the INSPIRE framework: To our knowledge, there are no published methods for learning modules and their dependencies that can handle variable discrepancy. We adapted the following five state-of-the-art methods that can run on a single dataset: <u>**GLasso**</u> - standard graphical lasso^24^, <u>**UGL**</u> - unknown group *L*_1_ regularization^25^, <u>**SLFA**</u> – the structured latent factor analysis^23^, <u>**WGCNA**</u> - weighted gene co-expression network analysis^8^, and <u>**MGL**</u> - module graphical lasso^11^ (see Methods for details). We adapted the input datasets such that we can apply these methods to datasets with variable discrepancy (Figure S1B):<u>**‘‐‐‐’**</u> - learning a model from only Dataset1 that contains all genes; <u>**‘Inter‐‐‐’**</u> - learning a model from the data on the overlapping genes (blue-shaded region in Figure 1) and assigning the rest of the genes to learned modules by using the *k*-nearest neighbor approach (i.e., based on the Euclidean distance between the gene’s expression and the expression of each of the modules), and <u>**‘Imp‐‐‐’**</u> – imputing missing values in Dataset2 and learning a model from the imputed data (see Methods for details on imputation) (Figure S1B). These adaptations lead to 13 competitors: 1) <u>GLasso1</u>, 2) <u>ImpGLasso</u>, 3) <u>UGL1</u>, 4) <u>ImpUGL</u>, 5) <u>WGCNA 1</u>, 6) <u>InterWGCNA</u>, 7) <u>ImpWGCNA</u>, 8) <u>SLFA1</u>, 9) <u>InterSLFA</u>, 10) <u>ImpSLFA</u>, 11) <u>MGL1</u>, 12) <u>InterMGL</u>, and 13) <u>ImpMGL</u>. In the experiments on synthetic data, we compared to all 13 methods, while in the experiments with two genome-wide ovarian cancer gene expression datasets which we will discuss in the subsequent sections, we only used the methods that are scalable (see Figure S9) These methods are indicated by the purple-shaded region in the table. The ‘Inter‐‐‐’ method is not applicable to GLasso and UGL, because GLasso and UGL learn a network of genes, not modules, and it is not obvious how to connect the genes that are present only in Dataset1 to the learned network. We do not consider an adaptation that applies the methods to Dataset2 only (‘‐‐‐2’). This is because, other than the genes in the overlap, Dataset2 has no genes (in the synthetic data experiments) or a very small number of genes (in the experiments with genome-wide expression data), which makes ‘‐‐‐2’ that uses only the samples from Dataset2 unlikely to outperform ‘Inter‐‐‐ ’ that uses all samples.

| Method | Description | Different ways to deal with missing data |  |  | Scalability<br>(see Figure S9) |
| --- | --- | --- | --- | --- | --- |
|  |  | ---1 | Inter--- | Imp--- |  |
| **GLasso** | standard graphical lasso<sup>24</sup> | GLasso1 | X | ImpGLasso | No |
| **UGL** | unknown group $L_1$ regularization<sup>25</sup> | UGL1 | X | ImpUGL | No |
| **SLFA** | structured latent factor analysis<sup>23</sup> | SLFA1 | InterSLFA | ImpSLFA | No |
| **WGCNA** | weighted gene co-expression network analysis<sup>8</sup> | WGCNA1 | InterWGCNA | ImpWGCNA | Yes |
| **MGL** | module graphical lasso<sup>11</sup> | MGL1 | InterMGL | ImpMGL | Yes |

#### Test log-likelihood

The *test log-likelihood* that measures how well the learned models fit unseen data is a widely used evaluation metric on probabilistic models^11,25,26^. We generated test data *Y* containing 100 samples, which was created in the same way as the training data *X* (see Methods). The 13 learned models are tested based on the same unseen data. Each method selects its own regularization parameter using the standard cross-validation (CV) test^27^ selecting *λ* with the best average CV test log-likelihood measured on Dataset1 in *X* (see Methods). We used the test set of Dataset1 to compute the test loglikelihoods for all methods since Dataset1 contains all genes. Figure 3A shows the average negative test log-likelihood per sample (lower the better) in (i) - (iv): (i) shows the methods that use only Dataset1, and (ii)-(iv) show Imp‐‐‐, Inter‐‐‐ and INSPIRE methods that use Dataset2 as well with varying numbers of genes in Dataset2 (Figure S1A). Each bar (except INSPIRE) displays a *p*-value from Wilcoxon signed rank test that measures how significantly INSPIRE is better than the corresponding method across 20 instantiations of the data (see Methods). The bars for the methods that use only Dataset1 display three - values, each for comparison to INSPIRE in (ii) - (iv). INSPIRE has significantly better test log-likelihoods than the methods that utilize one dataset (*p* ≤ 2.4 × 10^‒5^) and all the other 8 methods that can utilize multiple datasets (*p* ≤ 4.3 × 10^‒4^). This indicates that making use of multiple datasets by using INSPIRE has a great potential to increase the chance to infer the true underlying model. In (iv), ImpMGL, InterMGL and INSPIRE perform similarly as expected, and they are better than the other methods that utilize multiple datasets. The methods that utilize only Dataset1 (i) achieve worse average test log-likelihood than their multiple-dataset counterparts (ii)-(iv); and the test log-likelihood of most methods increase with the increasing number of overlapping variables, from (i) to (iv).

#### Module recovery

We then evaluated based on how well important aspects of the true underlying model are recovered by each method. We first checked whether pairs of genes that are assigned to the same module in the true model are in the same modules in the learned model. We used the rand index^28^ that measures how well pair of genes agree on being in the same or different modules between two models - the true model and a learned model. The rand index of 0 means that none of the genes agree on being in the same/different groups, while 1 means a perfect recovery of the modules. The evaluation based on module recovery is not applicable for GLasso1 and ImpGLasso, since they do not learn modules. As shown in Figure 3B, the module recovery performance of INSPIRE is significantly better than its 13 competitors. INSPIRE has significantly higher rand index than (i) the methods that utilize a single dataset (*p* ≤ 4.9 × 10^‒2^), and (ii)-(iv) the methods that use multiple datasets (*p* ≤ 6.6 × 10^‒2^).

#### Module dependencies

Then, we evaluated based on how well the inferred modules dependencies by each method are consistent with those in the true model. Since it is not clear how to map a module in the true model to the corresponding module in the learned model, we converted each module-based network model into the equivalent gene-based probabilistic model, using a well-established method^11^. It is not enough to get only high precision or recall, so we used the 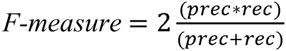 as an evaluation metric. As shown in Figure 3C, INSPIRE has the highest average F-measure that measures the accuracy of the dependencies learned by each method in all (i)–(iv). INSPIRE is significantly better than methods that utilize a single dataset (*p* ≤ 2.4 × 10^‒4^) and other methods that use multiple datasets (*p* ≤ 2.7 × 10^‒2^).

The methods that use only one dataset tend to have lower average rand-index (modules) and F-measure (module dependencies) than their multiple-dataset counterparts; and as the number of genes shared across datasets increases, the overall performance of the methods that utilize multiple datasets increases. This indicates that combining multiple datasets reveal underlying modules and their dependencies better, and INSPIRE is better than 13 alternative approaches in revealing underlying model.

### Evaluation on two genome-wide ovarian cancer expression datasets

Next, we evaluate INSPIRE based on the statistical robustness and biological relevance of the learned modules on two publicly available ovarian cancer gene expression datasets^29^ (Figure 4A): 1) OV1 that contains 18,113 genes and 28 patients (Affy U133 Plus 2.0 platform), and 2) OV2 that contains 8,331 genes in a total of 42 patients (Affy U95Av2 platform) (see Methods; Table S1).

We compared INSPIRE with six alternative methods that are scalable to genome-wide data (Table 1; Figure S9). The runtime of all the other methods when *p*=3, 000 is >10 hours, which means that running these methods on genome-wide data would be too slow to be used. 8,234 genes are presented in both datasets (rows in the blue-shared region in Figure 4A). As a preprocessing step, we standardized each dataset so that each gene has zero mean and unit variance across the samples within each dataset (See Methods). We used *k* = 91, where *k* is the number of modules, as selected by BIC on the *k*-means clustering applied to the imputed data matrix. We also present the results when *k* = 182 based on the biological plausibility of having on average 100 genes per module, in order to show that the outperformance of INSPIRE does not depend on one specific *k* value (Figure S2).

In the next three subsections, we show the results of the following evaluations (Figure 2B): 1) how well the INSPIRE model fits unseen data measured by test log-likelihood, 2) the statistical significance of the overlap between the learned modules (i.e., gene-module assignment) and known functional gene sets, and 3)how well the learned modules reflect putative regulatory relationships between transcription factors (TFs) and targets based on the ChEA database^30^.

### INSPIRE learns a statistically more robust low-dimensional representation model than alternative approaches

We first evaluated the learned low-dimensional representation (LDR) model based on the test-set log-likelihoods that measure how well the learned model can explain left-out test data in OV1 through the standard 5-fold cross-validation (CV) tests (see Methods). We used the test set of OV1 for computing the test log-likelihoods for all compared methods since OV1 contains almost all of the genes contained by either of the datasets. In Figure 4B, the best average test log-likelihood per sample across the tested λ values is plotted for each method. As can be seen in Figure 4B, INSPIRE achieves better test log-likelihood than six alternative methods, WGCNA1, InterWGCNA, ImpWGCNA, MGL1, InterMGL and ImpMGL (Table 1) for both *k* = 91 chosen by the BIC score (left panel), and *k* = 182, an alternative *k* value that results in modules with average size of 100 (right panel). Since MGL and INSPIRE may depend on the initialization of the model, the standard deviation across 10 runs of those methods with different initializations are represented by the error bars on the bottom panel in Figure 4B.

### INSPIRE modules are more significantly enriched for functional gene sets than alternative methods

INSPIRE uses a biologically intuitive low-dimensional representation (LDR) model for expression data, in which genes are assigned to *k* modules, and each module can be interpreted as biological processes performed by the genes in that module. Thus, whether each module is enriched for the genes that are known to be in the same functional categories can be a way to evaluate the biological relevance of the LDR inferred by INSPIRE. Here, we evaluated INSPIRE based on whether the learned modules are significantly enriched for known pathways from MSigDB^31^. We compared INSPIRE with six alternative methods, WGCNA1, InterWGCNA, ImpWGCNA, MGL1, InterMGL and ImpMGL (Table 1), using *k* = 91 chosen by the BIC score and *k* = 182, an alternative *k* value that results in modules with average size of 100. For each method, we chose *λ* that achieves the best CV test log-likelihood, a standard technique^27^.

We considered 1,077 GeneSets (pathways) from the C2 collection (curated gene sets from online pathway databases) of the current version of the MSigDB^31^ based on Reactome^32^, BioCarta and KEGG^33^. We excluded the pathways based on computational predictions from this collection. We computed the significance of the overlap between each GeneSet and each module measured by the Fisher’s exact test *p*-value, followed by the Bonferroni multiple hypothesis correction. Figure 4C and Figure S2A show the results of the functional enrichment analysis for *k* = 91 (chosen based on BIC) and *k* = 182, respectively. In each scatter plot, a larger portion of the dots lie above the diagonal, which implies that the INSPIRE modules are more significantly enriched for known pathways than those inferred by the alternative approaches. This indicates that the INSPIRE is better at identifying biologically coherent modules based on prior knowledge more accurately than the alternative methods.

### INSPIRE modules are more significantly enriched for putative targets of the same transcription factor than alternative approaches

As an alternative way to evaluate the biological coherence of the learned modules, we checked how significantly the modules are enriched for the genes that have been shown to be bound by the same transcription factors (TFs). The ChEA database^30^ provides a large collection of TF-target interactions captured in previously published ChIP-chip, ChIP-seq, ChIP-PET and DamID (referred herein as ChIP-X) data. For each of 107 TFs in the ChEA database^30^, we computed the significance of the overlap between each module and each TF’s putative targets from ChEA database measured by the Fisher’s exact test *p*-value followed by the Bonferroni correction. Figure 4D and Figure S2B show the results of our ChEA enrichment analysis for *k* = 91 (chosen based on BIC) and *k* = 182, respectively. In each scatter plot, a much larger portion of the dots lie above the diagonal, which indicates that INSIRE modules are biologically more coherent, i.e., more significantly enriched for putative targets of the same TF. In Figure 4D and Figure S2B, we indicate with a blue dot a TF that resides in the same module as the module that is enriched for the TF’s putative targets. We do not expect all dots to be blue-colored (i.e., all TFs being in the same modules as their putative targets), because the protein level of TF may not be correlated with its mRNA expression level. It is still interesting to see that INSPIRE modules are more significantly enriched for the genes that have been shown to be bound by the same TFs in ChIP-X data.

### Application to nine genome-wide ovarian cancer expression datasets

Encouraged by the in-depth evaluation described above, we applied INSPIRE to nine expression datasets that comprise 1,498 ovarian cancer patient samples downloaded from the TCGA project website and the Gene Expression Omnibus (GEO)^34^ (Figure 2C). This corpus of data consists of publically available transcriptomic characterizations of ovarian cancer across nine distinct studies where gene expression data collected in different studies come from distinct platforms. This data is therefore a perfect corpus to apply the INSPIRE method for a variety of reasons. First, there is a sufficient sample size across studies to resolve distinct modules that are robust across datasets. Second, our method will outperform more naive approaches by imputing missing genes through leveraging shared structure across the data, and will therefore increase the resolution to detect robust modules. Finally, there are known subtypes in ovarian cancer as identified by the TCGA ovarian cancer study^13^, and we anticipate that our approach will not only re-identify these subtypes based on the expression of our inferred modules, but will also further resolve potential molecular drivers of these subtypes through ancillary analyses of the INSPIRE inferred modules. These ancillary analyses are described below. We repeated our analyses for this application using varying module counts *k* = {90,129,181} thatcorrespond to the average number of 200, 140 and 100 genes respectively in each module and for varying sparsity tuning parameters *λ* = {0.01,0.03,0.1}; and we observed that all results were highly robust for the varying values of *k* and *λ*. We reported results from our biological analysis for, as selected by BIC for the *k*-means clustering applied to the imputed data matrix, and *λ* = 0.1 which leads to the sparsest network of modules, given that sparsity is of key importance in learning and the interpretation of a high-dimensional conditional dependence network.

We evaluated the learned low-dimensional representation (LDR) consisting of 90 modules and the corresponding latent variables based using three evaluation metrics:

1) We performed gene set enrichment analysis to characterize each module based on its associated genes (see Table S3 for the gene set enrichment analysis results together with the significance).
2) We analyzed the associations between the learned latent variables, each representing a module, and six important phenotypes in cancer, including resectability which was defined by the residual tumor size after surgery, survival, and four histopathological phenotypes manually curated based on the histopathology in the TCGA ovarian cancer data (see Table S3), and we used inferred INSPIRE latent variables as features for predicting those phenotypes. Figure 5A shows the association between the learned latent variables with the six important phenotypes, and Figure 5B compares INSPIRE to the following based on the prediction of those phenotypes: i) principal component analysis (PCA)^20^ – an unsupervised LDR method; ii) subnetwork analysis^15^ – a supervised LDR method; and iii) all genes when no LDR is learned. The histopathological phenotypes are provided as a resource for this paper (Table S4), and residual tumor size and survival are available on the TCGA web site.
3) We used the inferred latent variables to identify new subtype definitions in ovarian cancer. We compared INSPIRE subtypes to i) the subtypes recently described by the TCGA ovarian cancer study^13^, and ii) the subtypes learned by a method that uses mutation profiles for the network-based stratification of cancer patients (NBS)^35^, based on how relevant they are to genomic abnormalities in ovarian cancer. Detailed information concerning expression datasets used in the INSPIRE analysis is presented in Table S2,and the processing of the expression data is described in Methods.
4) We perform both statistical and biological experiments to show that *HOPX* is a potential molecular driver from tumor-associated stroma in a module that differentiates the patients with increased percent stroma, infiltrative stroma, and desmoplastic stroma.

**Figure 5.**
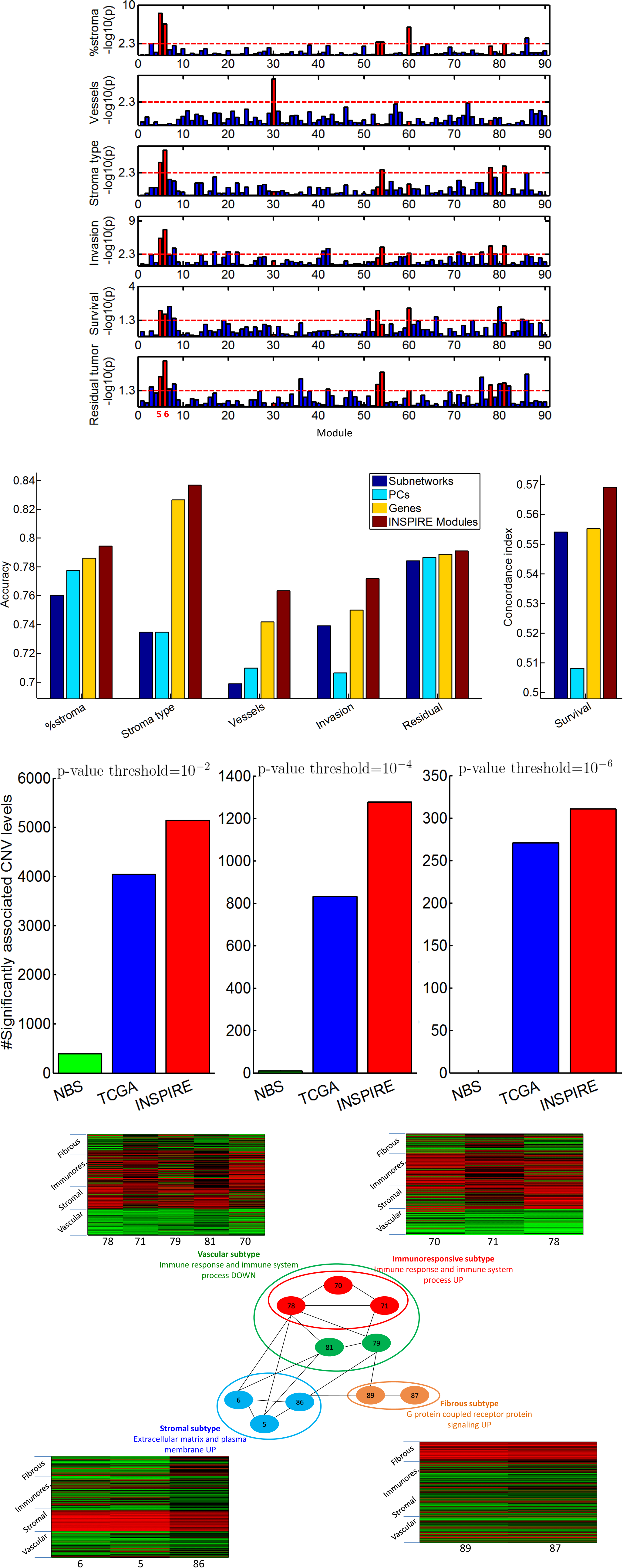
**(A)** For each of 90 INSPIRE modules (x-axis), the – log_10_ *p* from the Pearson’s correlation is shown (y-axis) for six different histological and clinical phenotypes. The *p*-value threshold (shown by red dotted horizontal lines) is 5 × 10^‒3^ for histological phenotypes, and 5 × 10^‒2^ for clinical phenotypes, which are harder to predict. We highlight modules 5, 6, 53, 54, 60, 78 and 81 that are significantly correlated with at least three of the six phenotypes by red. We also highlight module 30 by red since it is the only module that has a significant correlation with the vessel formation phenotype. Modules 5 and 6 achieve the first or second rank in terms of the significance of correlation with five of the six phenotypes. **(B)** For four different methods (the subnetwork markers, principal components, all genes and INSPIRE latent variables, the prediction performance is compared for six prediction tasks in cross-validation setting. **(C)** For three different Pearson’s correlation *p*-value thresholds (10^‒2^,10^‒4^,10^‒6^ respectively from left to right), the number of CNV levels that are significantly associated with the learned subtypes are shown for two published methods and INSPIRE. **(D)** The modules that differentiate the subtypes that are learned using INSPIRE features and the interactions among those modules as learned by INSPIRE.The modules are grouped and colored according to the subtypes they differentiate. Next to each one of the four module groups, there is the heat map of the features corresponding to the modules in this module group.

### Negatively correlated modules show distinct pathways and potential regulatory TFs enrichment

We emphasize that the key goal of INSPIRE is to reduce the dimensionality of expression data in a biologically intuitive way and in such a way as to capture important dependencies. Given that the gene regulatory network is known to be highly modular^36^ and dimensionality reduction is the key goal, we chose to focus on module-level dependencies rather than gene-level dependencies. The ability to capture the high-level abstraction of the dependencies among gene expression levels is a key goal and advantage of INSPIRE. As a result of the INSPIRE model assumptions, expression of genes in the same INSPIRE module would tend to be positively correlated, and positive correlation in expression levels across patients is an important property - expression activated or deactivated within similar sets of patients. Genes with strong negative correlations are likely to be highly related functionally, however they would have completely different regulatory mechanisms (e.g., different transcription factor binding) and biological interpretation. In Figure S8, we show scatter plots in which each dot corresponds to a GeneSet (from the pathway databases or transcription factor (TF) binding information) and we plot the maximum ‒log10(p) obtained by each model (axis).

Figure S8A (top) demonstrates that the modules that are strongly negatively correlated with each other show very distinct pathway (left) enrichment as well as TF binding enrichment (right). In Table S9, the significance of enrichment from five negatively correlated module pairs with the biggest absolute correlation listed for five pathways or transcription factors for which the highest enrichment difference between the negatively correlated modules is observed.

We also compared between following two models in terms of functional enrichment of the modules: I) two negatively correlated modules are defined as two separate modules as in the original work; and II) instead of the two negatively correlated module, there is one hypothetical module that contains all genes in the two negatively correlated modules. Figure S8A (bottom) compares between model I (y-axis) and model II (x-axis) in terms of functional coherence based on the pathway database (left) and putative TF binding targets (right). Model I reveals more functionally coherent modules than model II, which justifies our modeling assumption that negatively correlated genes need to be in separate modules.

### INSPIRE latent variables are significantly associated with clinical and histologic phenotypes in cancer

To gain relevant biological insight from ovarian cancer (OV) transcriptome data, we used the 90 inferred latent variables from the INSPIRE model as a lower dimensional representation (LDR) of transcriptomic profiles across patients (Figure 2C), that captures robust cross dataset patterns of gene expression. We evaluated the clinical relevance of these latent variables by measuring the statistical association between these latent variables and histopathological phenotypes of tumor. The morphological interpretation of histologic sections of tumor forms the basis of diagnosis, aggressiveness assessment, and prognosis prediction. Pathologists examine the tumor diagnostic images based on semi-quantitative histologic phenotypes of the tumor such as invasion pattern and percent stroma to predict the aggressiveness of cancer. Identifying the molecular basis for these histologic phenotypes will advance the understanding of the molecular biology of ovarian cancer. We manually examined five histologic phenotypes for 98 randomly selected patient images from TCGA: percent stroma, percent tumor, vessel formation, stroma type, and invasion pattern (details in Methods; Table S4). For each pair of a histologic phenotype and a latent variable from the INSPIRE model, we performed the Pearson’s correlation test that produces a correlation coefficient and a *p*-value. Table S3 lists the *p*-values from these association tests of INSPIRE latent variables, with each of the five histologic phenotypes. Figure 5A shows the correlation of each latent variable with each of the histologic phenotypes. Since percent stroma and percent tumor phenotypes are almost perfectly (anti-) correlated, we only included percent stroma in Figure 5A. We used *p*-values from a likelihood ratio test for a Cox proportional hazards model to determine the significance of association of a gene with patient survival, and we used-values from the Pearson’s correlation test for tumor resectability.

Modules 5 and 6 show high correlations with the histopathological phenotypes, such as percent stroma, stroma type and invasion pattern. As shown in Figure 5A, those modules are also associated with patient survival and tumor resectability. We observed that the quantity of residual tumor after surgery is positively correlated with the amount of tumor-associated stroma, where increased residual tumor, i.e. low resectability, is an important and a previously known indicator of poor patient prognosis. Although the latent variables of module 5 and module 6 show high expression correlation (the correlation coefficient between the module 5 latent variable and the module 6 latent variable is 0.84), these two modules are functionally fairly different. Figure S8B compares module 5 and module 6 in terms of the pathways and putative TF targets that are enriched in these modules. There are handful of dots that are distant from the diagonal line implying that module 5 and module 6 exhibit several unique biological properties. In Table S10, the significance of enrichment from module 5 and 6 are listed for five pathways or transcription factors for which the highest enrichment difference between the modules is observed.

To examine the difference between module 5 and module 6 in terms of phenotypes associated with them, we compared between the following two models in an experiment where the latent variables are used as *features* in predicting six different phenotypes (percent stroma, stroma type, vessel formation, invasion pattern, resectability and survival): I) module 5 and module 6 exist as two separate modules as in the original work; and II) instead of module 5 and module 6, there is one hypothetical module that contains all genes in modules 5 and 6. As shown in Table S11, we observed that module 5 and module 6 are significantly predictive of distinct sets of phenotypes, and interestingly, either module 5 or module 6 is always better in terms of predictability of phenotypes than the hypothetical module containing all genes in modules 5 and 6, which means model I is a better predictor of all six phenotypes than model II. Thus, even if modules 5 and 6 are highly correlated with each other, the genes in these modules need to be separated into the two modules.

### INSPIRE latent variables are more predictive of clinical and histologic phenotypes in cancer than other kinds of LDRs and all genes

Many biological processes are performed by a group of genes rather than individual genes, and as a result, many *complex* phenotypes and clinical outcomes can be explained based on module activity levels rather than individual genes. Moreover, expression level of an individual gene is often noisy and even if it was not, it still may not be perfectly correlated with a protein level of a true regulator for a phenotype.

To test this hypothesis and further demonstrate the effectiveness of INSPIRE as an LDR of gene expression data, we used the INSPIRE latent variables as *features* in prediction tasks, and we compared INSPIRE with the following methods: i) principal component analysis (PCA)^20^ – the most widely used unsupervised LDR method; ii) subnetwork analysis^15^ – a powerful supervised LDR method that extracts network markers; and iii) all genes when no LDR is learned. The subnetwork analysis method^15^ learns small sub-networks of genes in a given large PPI network, based on expression data and a particular prediction task. For example, for a stroma type prediction (fibroblast/desmoplastic), it learns subnetworks of genes in a given PPI network such that the average expression level of each sub-network significantly differentiates the two patient groups based on the classes of stroma type. This method is a supervised method in that the subnetworks are learned such that they can explain a particular phenotype well. On the other hand, INSPIRE is an unsupervised method in that the result does not depend on a particular prediction task. Each of INSPIRE latent variables, subnetworks, principal components, and all genes is considered as a set of features in predicting six different phenotypes; percent stroma, stroma type, vessel formation, invasion pattern, resectability and survival (see Methods for details). The result of the comparison shows that the features learned by INSPIRE show the best prediction performance measured among all methods considered (Figure 5B). This result strengthens our claim that the INSPIRE latent variables provide informative lower-dimensional features for prediction tasks.

Because INSPIRE groups genes in multiple datasets into a set of modules, most modules may include a significant number of genes whose expression is not correlated with the predicted phenotype. In order to examine the effect of those genes in phenotype prediction tasks, we generated four hypothetical module sets by excluding 20%, 40%, 60%, and 80% of the genes whose expression levels in training samples are least significantly associated with the respective phenotype from each of 90 modules, and repeated the phenotype prediction experiments for those four hypothetical module sets. Table S12 shows that the original INSPIRE latent variables which correspond to the module set including non-discriminative genes perform the best and in most cases, the performance even decreases when top 20% of the most discriminative genes are left. This result indicates that latent variables resulting from the contribution of all genes make robust features informative of the phenotypes.

### Subtypes inferred based on INSPIRE latent variables are highly relevant to genomic abnormalities in ovarian cancer

Cancer is a heterogeneous disease with multiple distinct genetic drivers, where identifying subtypes of cancer relevant to potential genetic drivers is a primary goal of the field of cancer biology. Here, we cluster ovarian cancer patients from the TCGA study^13^ (560 samples) into four subtypes by using the latent variables learned by the INSPIRE method as *features for clustering* patients (details in Methods). Table S7 lists the assignment of the patients in the TCGA ovarian cancer data to the four INSPIRE subtypes.

To examine the relevance of the INSPIRE-based subtypes to the potential drivers of ovarian tumor, we checked the significance of the association between the subtypes with copy-number variation (CNV) of genes, an important genomic abnormality that can drive cancer (Figure 5C and Figure S3A). We focused on CNV for this test instead of mutation since ovarian cancer has been characterized as a c-class cancer (as opposed to m-class, where ‘m’ represents mutation) in which CNV is more prevalent than mutations^37^. For each CNV (as quantified by the CNV level), we performed a multivariate linear regression using the INSPIRE subtypes, where we computed a *p*-value (from the regression *f*-statistic) to ascertain how well the INSPIRE subtype regression model fits a given CNV. We then compared the number of CNVs with significant INSPIRE *p*-values (determined by varying thresholds; see Figure 5C) to the number of CNVs with significant *p*-values from the following two approaches: 1) the subtypes learned by using a method that uses mutation profiles for the network-based stratification (NBS) of cancer patients^35^, and 2) the subtypes inferred from a recent TCGA ovarian cancer study^13^. Figure 5C shows that INSPIRE results in subtypes that are more associated with CNV based genomic abnormalities than alternative approaches. In Figure S3A, we show the comparison for varying numbers of modules (*k*), for varying sparsity tuning parameters (*λ*), and for varying *p*-value thresholds, which shows that the results are robust to varying hyper-parameters. Figure 5C and Figure S3A indicate that INSPIRE further resolves subtypes as defined by the potential genomic drivers of ovarian cancer when compared to alternative approaches. In the Supplementary Note 1, we list the CNV levels that are significantly correlated with each of the four subtypes. The enrichments of those CNV levels with the MSigDB^31^ C2 (curated gene sets) categories and the corresponding – log_10_ *p* are also listed for each subtype.

### Subtypes revealed by INSPIRE and their relationships with the TCGA subtypes

Figure 5D reveals a sub-network learned by modules from an INSPIRE model using parameters *λ* = .1 and *k* = 90 (chosen based on BIC). This subnetwork contains modules that are differentially expressed in one of the four subtypes, as represented by the heatmaps in Figure 5D. The differentially expressed modules, termed *marker modules*, are determined for each subtype by comparing the subtype versus the other three subtypes, using the Significance Analysis of Microarrays (SAM) algorithm^38^ implemented in the R package *siggenes*. Table S8 lists the enrichment of the marker modules with the MSigDB^31^ C5 (GO gene sets) and the corresponding – log_10_ *p*. We observed that the set of marker modules (Table S8) have a significant overlap (*p* = 2.4 × 1CT^3^) with the set of modules that have significant associations with at least three of the six phenotypes (the modules colored in red in Figure 5A and Table S3 except module 30). Not surprisingly, the INSPIRE subtypes show diverse histologic features across subtypes, and we accordingly termed the INSPIRE subtypes ‘vascular’, ‘stromal’, ‘immunoresponsive’, and ‘fibrous’. See Figure 5D and Table S8, where the marker modules for the vascular, stromal, immunoresponsive, and fibrous subtypes are colored in green, blue, red and orange, respectively.

Table S5 shows a confusion matrix that describes the overlap between the INSPIRE subtype assignments and the TCGA subtype assignments^13^ together with the *p*-values for the significance of the overlap for the highly-overlapping subtypes. There is a more significant overlap for the vascular-proliferative pairs and stromal-mesenchymal pairs, which implies that the proliferative-like and mesenchymal-like subtypes are highly conserved across different OV datasets, which is consistent with the finding of Way et al.^39^. Although the INSPIRE subtypes have a statistically significant overlap with the TCGA subtypes, the INSPIRE subtypes show much stronger association with genomic abnormalities, as mentioned above (see Figure 5C). We further include the description of the stromal subtype here since it is characterized by the high expression of modules 5 and 6, which are strongly associated with the six important phenotypes in cancer (Figure 5A). See the Supplementary Note 2 for the characterization of the other three (‘vascular’, ‘immunoresponsive’ and ‘fibrous’) subtypes.

The stromal subtype is characterized by high expression of modules 5, 6 and 86 (Figure 5D) and associated increased percent stroma, infiltrative growth pattern, and desmoplastic stroma (Figure S3B (i), (ii), (iii)). Modules 5 and 6 are significantly enriched for proteinaceous extracellular matrix gene sets (Table S8), which is likely due to increased percent stroma. In Figure 5D, there are quite a few edges between modules associated with the vascular subtype and those associated with stromal subtype, which suggests a strong association between the increased stromal components and neovascularization of the tumor. This likely reflects the known tumor neovascular niche in cancer that involves proangiogenic factors release from tumor stroma along with the vasculature itself^40^. This is supported by multipotent mesenchymal stromal cells having unique immunoregulatory and regenerative properties^41^. A substantial amount of the tumor stroma is composed of immune cells, and the net effect of the interactions between these various immune cell types and the stroma participates in determining anti-tumor immunity and neovascularization potential^42^. We note that the immune system modules 78 and 81 that are connected to extracellular matrix modules 5 and 6 are also up-regulated in the stromal subtype (Figure 5D). Stromal subtype is a significant predictor of poor patient survival (Cox proportional hazards model log-rank *p* = 8.8 × 1CT^2^) with a median survival of 914 days. Cancers associated with a reactive stroma is typically diagnostic of poor prognosis^43^, and we observed that median survival of the stromal subtype is the smallest among all subtypes. Stromal subtype has a significant overlap (*p* = 1.03 × 10^‒35^) with the mesenchymal subtype discovered by TCGA^13^ (Table S5).

### INSPIRE provides novel insights into molecular basis for ovarian tumor resectability

Riester et al. identified POSTN as a candidate marker for tumor resectability in ovarian cancer^44^, where the resectability phenotype was defined by the residual tumor size after surgery. The authors showed that high POSTN expression is strongly associated with poor tumor resectability, even more so than a multigene model chosen by leave one out cross-validation across 1,061 samples in 8 datasets including the TCGA^13^ and Tothill^45^ datasets. POSTN is a member of module 6 that shows the most significant association with resectability among all 90 modules (Figure 5A). We, therefore, compared our supervised prediction model using the INSPIRE latent variables corresponding to modules 5 and/or 6 to a model that contains just POSTN to determine whether the genes in module 5 and the genes in module 6 other than POSTN provide any information to the prediction of resectability in addition to the information provided by the POSTN expression. We observed that when training on TCGA data^13^ including the clinical covariates, the models trained using 1) modules 5 and 6 together, 2) module 6 only, 3) module 5 and POSTN together, and 4) module 5 only, outperformed the model with the known marker for resectability, POSTN, when tested in the Tothill^45^ dataset (see AUC values in Figure 7). TCGA data^13^ was used for training because of its large sample size. Tothill^45^ was used for testing, because it has the largest sample size except TCGA data (Table S2) and contains the most fine-grained information on the residual tumor size. Additionally, the proportion of optimally and sub-optimally debulked patients was similar between TCGA and Tothill data. We used a stringent definition of resectability (0 cm vs. >0 cm) (see Methods).

**Figure 7.**
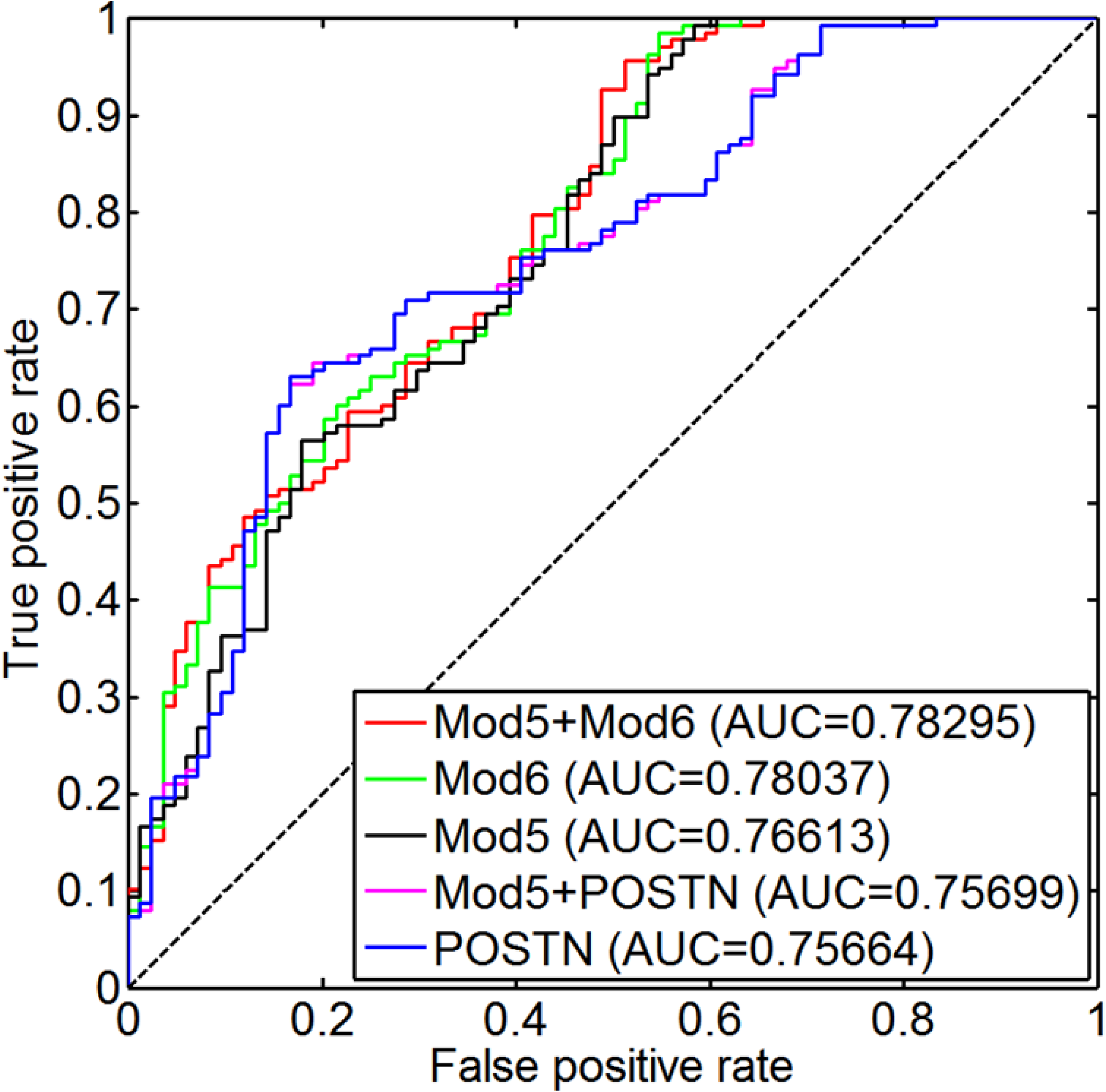
ROC curve of the supervised models for resectability prediction trained in TCGA and tested in Tothill data. Different combinations of POSTN and the INSPIRE features corresponding to modules 5 and 6 are used for training each model. The clinical covariates age and stage are also included in all models. AUC of each model is shown in the legend.

Since module 6 contains POSTN, outperformance of 1)-3) means that the modules 5 and 6 representing the expression of genes in module 5 and/or module 6, which are significantly predictive of stromal histology features and resectability, add information to the prediction of resectability by POSTN in a cross-dataset analysis. Outperformance of 4) means that the module 5 representing the gene expression levels in module 5, which does not contain POSTN, is a better predictor of resectability than POSTN. When we repeated this experiment with no clinical covariates (age and stage) in the training, the models including module 6 outperformed the model that includes only POSTN, which means the genes in module 6 other than POSTN add information to the prediction of resectability by POSTN (see AUC values in Figure S5). The modules 5 and 6 with strong stromal and mesenchymal properties (see below) provide potential novel molecular basis for tumor resectability.

### INSPIRE modules and the conditional dependence network among them

Here, we discuss the modules that show significant correlations with many of the histological and clinical phenotypes in the TCGA ovarian cancer data or that achieve the only significant correlation with a phenotype among all modules (see Figure 5A and Table S3).

Module 5 contains known EMT inducers *ZEB1, SNAI2* and *TCF4 (E2.2)*^46^ as well as multiple other genes known to be important in focal adhesion^47^, extracellular matrix interaction^48^, extracellular matrix organization^49^, and markers of cancer-associated fibroblasts (*PDGFRB, PDGFRA*)^50^ (see Table S3). Similarly, module 6 contains EMT inducer *TWIST1*^46^, many extracellular matrix genes, as well as genes associated with senescence and autophagy, collagen genes, and the well validated predictor of tumor resectability, *POSTN*^44^ (see Table S3). These two modules are prime candidates for genes driving EMT associated tumor aggression. Although modules 5 and 6 have many shared GO categories and pathways, they are likely to represent fairly different biological processes (Figure S8B). When we combined these two modules and used one latent variable that represents the two modules, the overall prediction results became worse (Table S11).

While modules 5 and 6 contain known drivers of EMT and extracellular matrix genes and these modules are also associated with tumor-associated stroma/mesenchymal phenotypes, we found other modules with significant correlations with most the histological and clinical phenotypes. Additionally, active area of research in cancer biology is to identify pathways and genes driving tumor aggression. This includes genes associated with cancer stem cells (i.e., tumor-initiating cells)^51–54^. Module 78 contains genes indicative of hematopoietic cell lineages likely because it includes many innate immune response genes, as well as multiple innate immune response signaling pathways including cytokine cytokine receptors, toll like receptors, and TCR signaling. Module 78 also contains a known EMT inducer ZEB2^46^. This indicates that module 78 may capture aspects of tumor associated inflammation, a known contributing factor to EMT^55^. Module 81 includes genes that regulate the MAPK and ERK cascades, signal transduction pathways that are known to be upstream of multiple oncogenic process^56^. Module 54 represents genes involved in pro-apoptotic and cell cycle regulation. GADD45 genes, known to be upstream of JNK signaling^57^, are present along with JUN and FOS. In addition, this module contains KLF4 and KLF6, which like GADD45, are known to repress cell cycle arrest and associated cyclin-dependent kinase inhibitors^58^. Modules 30 and 54 are indicative of the likely metabolic shift that cancers cells undergo as these modules are enriched in metabolic and biosynthesis pathways. When considering these modules jointly, we get a picture of multiple processes (Figure S6), and potential tumor cell subpopulations, that populate the tumor microenvironment and perpetuate aggressive tumor states in subpopulations of patients.

One of the advantages of the INSPIRE framework over naïve clustering algorithms is that it suggests potentially biologically relevant interactions or couplings between the modules. These interactions can be used to motivate higher-level hypotheses about the coupling of disease specific processes.

### INSPIRE reveals a previously unknown stroma-associated marker *HOPX*

Given the association of the genes in the modules 5 and 6 with aggressive stroma and patient prognosis, and the significance of the modules 5 and 6 in differentiating the stromal subtype, we were interested in understanding if the modules 5 and 6 capture a prognostic signature that generalizes across other cancers. Prognostic genes are more likely to be shared by distinct tumor types than would be expected by random chance likely because of prognostic mechanisms that generalize across cancers (e.g. metastatic potential or immune system evasion), and conversely, cancer-specific prognostic genes are less frequent than would be expected by random chance^59^. Therefore, to further annotate modules 5 and 6, we performed a pan-cancer analysis to check whether the genes contained in those modules are significantly associated with survival in six publicly available datasets^6,45,54,60–62^ from five cancer types: ovarian cancer, breast cancer, acute myeloid leukemia, glioblastoma and lung cancer (see Table S6 for the details of these datasets). We used *p*-values from the likelihood ratio test for a Cox proportional hazards model to determine the significance of association of a gene with patient survival, and we considered a *p*-value ≤ 0.05 to be significant. We observed that the genes in module 5 and the genes in module 6 are significantly associated with survival in at least three of the six datasets (Fisher’s test statistic *p*-value = 1.68 × 10^‒8^ for module 5 and = 4.44 × 10^‒8^ for module 6). For breast cancer, we used the Osloval (the test data) but not Metabric (the training data with 1981 samples from the same study^6^ with Osloval) for breast cancer because we need the sample sizes to be similar across datasets such that the meta-analysis is not dominated by a single cancer type.

To further investigate the specific genes that are associated with patient survival across cancer types in these modules, we computed a combined *p*-value statistic using Fisher’s combined probability test for the association of each gene with patient survival in a meta-analysis of the six datasets from these five cancer types. *HOPX*, which is in module 5 achieved the lowest combined *p*-value among all genes in module 5 or module 6, and the third lowest combined *p*-value genome-wide (*p* = 1.32 × 10^‒10^).Thetoptwo genes that yield smaller *p*-values than HOPX genome-wide are CD109 (*p* = 2.49 × 10^‒11^) and SKAP2 (*p* = 3.55 × 10^‒11^) neither of which is in the module 5 or module 6 (Figure 6A). As shown in the previous sections, module 5 (containing 183 genes) is highly associated with percent stroma (Figure 5A), and is significantly enriched (*p* = 8 × 10^‒5^) for the known drivers of EMT that has been shown to contribute to poor patient survival. Not all 183 genes in module 5 would play a key role in the formation of tumor-associated stroma or EMT, and in fact, many of the genes in module 5 would simply have correlated expression pattern with key genes in these processes. We hypothesize that such genes have robust association with survival enough to be conserved across different cancer types, given the previously known association between tumor-associated stroma and patient survival. We note that known EMT drivers *ZEB1, SNAI2* and *TCF4* in module 5 have significant associations with survival in our pancancer analysis (*p*-values 8.5 × 10^‒6^,5.3 × 10^‒4^,1.3 × 10^‒3^ and rankings 153, 749 and 1098 respectively out of 11119 total genes). Thus, our pan-cancer analysis that highlights *HOPX* in module 5 led to us to consider HOPX as a potential molecular marker strongly associated with percent stroma and tumor aggression. Additionally, HOPX is one of the 15 genes in module 5 (out of 183 genes) that have been classified as ‘candidate regulators’^63^. Gentles et al. have defined a list of about 3,000 genes as candidate regulators, those that have a potential regulatory role in the broad sense (not specific to cancer): transcription factors, signaling proteins and translational initiation factors that may have transcriptional impact^63^. This implies that HOPX could be a regulator in the stroma-associated processes.

**Figure 6.**
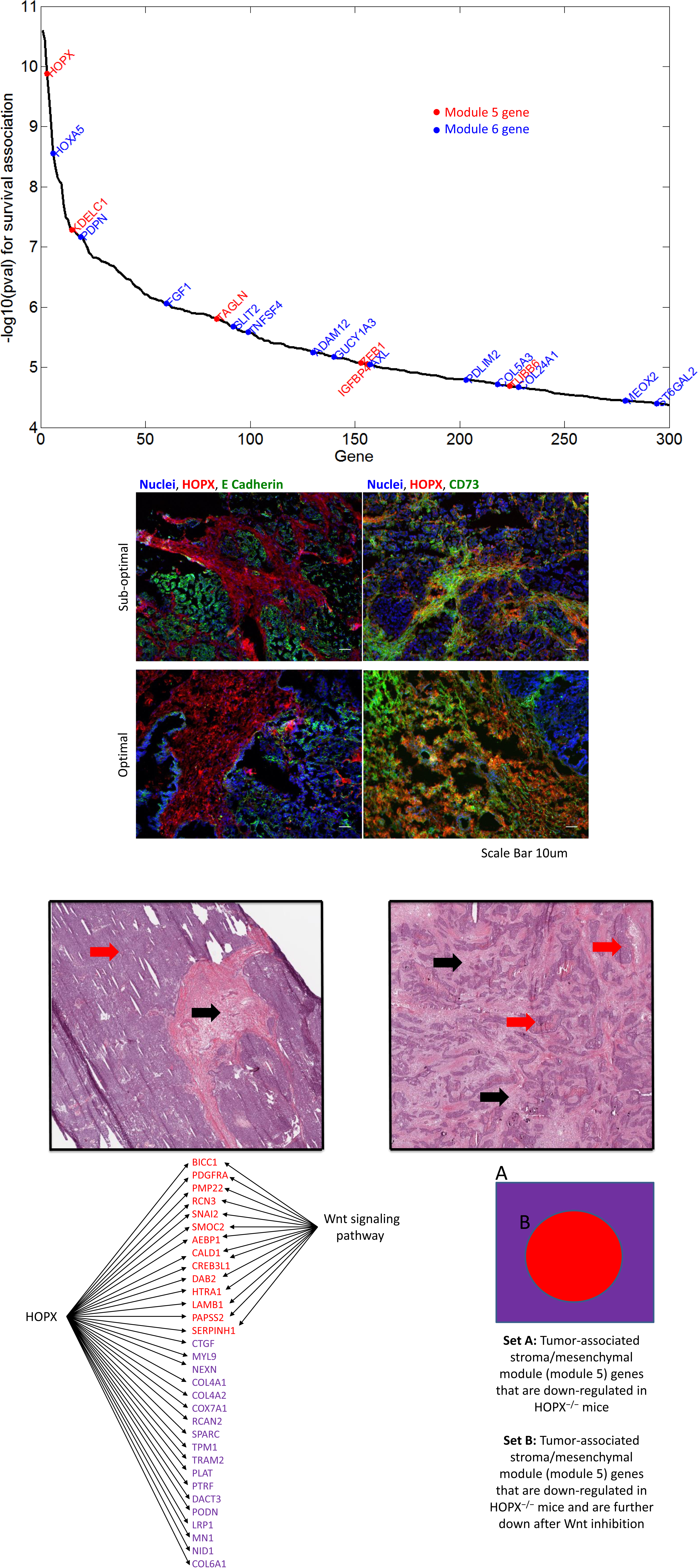
**(A)** Fisher’s combined p-values for survival (y-axis) are shown for the top 300 genes (x-axis) which achieve the most significant survival association in the pan-cancer survival analysis. Module 5 genes are shown by red and module 6 genes are shown by blue. **(B)** Fluorescent staining of ovarian tumors from sub-optimally bulked and optimally debulked patients. Each row is a single patient. HOPX is localized to the stroma and does not overlap with *ϵ* Cadherin positive cancer cells. HOPX does however overlap with CD73, a mesenchymal stem cell marker **(C) Left:** Expansile growth pattern of high-grade serous carcinoma associated with optimal resectability and low HOPX expression from the TCGA ovarian cancer study. Note high percentage of carcinoma (red arrow) and low percentage of stroma (black arrow). Hematoxylin and Eosin, 100X. *Right:* Infiltrative growth pattern of high grade serous carcinoma associated with low resectability and high HOPX expression from the TCGA ovarian cancer study. Note high percentage of stroma (black arrows) compared with carcinoma (red arrows). Hematoxylin and Eosin, 100X. *(D)* A total of 32 genes that are potential targets of HOPX are shown. The purple-colored genes are the targets whose expression does not depend on Wnt signaling, and the red-colored genes are the targets of HOPX which are down-regulated in HOPX^-/-^, and further down upon Wnt inhibition in HOPX^-/-^. It is highly likely that the expression of the red-colored genes are driven by both HOPX and Wnt signaling pathway.

### *HOPX* is a putative driver for the tumor-associated stroma/mesenchymal module (module 5)

*HOPX* is an unusual HOX protein that does not contain a DNA binding domain, and has been implicated in multiple aspects of cardiac and skeletal muscle development through recruitment of histone deacetylases^64–66^. It has been suggested to have tumor suppressive function in other cancer types^67–69^, which confounds how its expression in OV is associated with several poor outcomes. This may also reflect different roles for *HOPX* in ovarian tumor-associated stromal tissue.

Previous studies characterize *HOPX* as a mediator of canonical Wnt and Bmp signaling, and may play key roles in maintaining a stem cell like state^70^. In our further analysis of HOPX, we observed that *HOPX* is one of the top candidate expression regulators for ovarian cancer^63,71^. To understand how HOPX is associated with the genes in the tumor-associated stroma/mesenchymal module (module 5), we compared these genes with those down-regulated in HOPX^-/-^ mice compared to HOPX^+/-^ control mice^70^ and found a significant enrichment based on the Fisher’s exact test (*p* = 1.5 × 10^‒3^). Those two results together suggest HOPX is a good candidate driver for the tumor-associated stroma/mesenchymal module, as many of the other genes in module are putative downstream targets of HOPX, either directly or indirectly. Figure S4C shows the enrichment *p*-value and the fold enrichment of the genes in the tumor-associated stroma/mesenchymal module with the down-regulated genes in HOPX^-/-^ mice for varying fold change of expression of the down-regulated genes (x-axis).

Furthermore, genes down-regulated in HOPX^-/-^ mice after addition of XAV939, a potent inhibitor of Wnt signaling to HOPX^-/-^ mice^70^, are even more significantly enriched (*p* = 1 × 10^‒8^) for genes in the tumor-associated stroma/mesenchymal module. The HOPX protein is a potent Wnt inhibitor ^70^, therefore in the HOPX^-/-^ mice Wnt is activated, and genes inhibited by Wnt are also turned off. When the Wnt inhibitor is applied to the HOPX^-/-^ mice the genes inhibited by Wnt are no longer turned off, and the down-regulated genes are more specific to genes specifically activated by HOPX, instead of being a mixture of genes activated by HOPX and inhibited by Wnt. In addition, it is not surprising to see a higher enrichment upon Wnt inhibition, because canonical Wnt signaling has been implicated in the regulation of the stromal activity of MSCs^72,73^. Figure S4D shows the enrichment *p*-value and the fold enrichment of the stroma/mesenchymal module genes that are down-regulated in Wnt-inhibited HOPX^-/-^ mice for varying fold change of expression of the down-regulated genes (x-axis).

These results suggest that the genes in the tumor-associated stroma/mesenchymal module which are down-regulated in both HOPX^-/-^ mice and Wnt-inhibited HOPX^-/-^ mice are good candidates as downstream targets of HOPX. Figure 6D shows those 32 potential targets of HOPX. The purple-colored genes are the targets that are down-regulated in HOPX^-/-^, and their expression does not change significantly (|*FC* change| ≤ 0.55) upon Wnt inhibition. On the other hand, the red-colored genes are the targets of HOPX which are down-regulated in HOPX^-/-^, and they are down-regulated further upon Wnt inhibition (|*FC* change| ≥ 0.93). It is highly likely that the expression of the red-colored genes in Figure 6D are driven by both HOPX and Wnt signaling pathway. We note that *HOPX* is, therefore, a potential driver for *SNAI2*, which is involved in EMT^74^ and *AEBP1*, which is a stromal adipocyte enhancer-binding protein.

### *HOPX* is a molecular marker of aggressive tumor stroma

To further disentangle the molecular underpinnings of the tumor-associated stroma/mesenchymal module, we stained tumor sections with antibodies against HOPX. We co-stained with E cadherin, a tumor epithelial cell marker. Patient samples were selected based on patient survival and optimal debulking (see Methods for details). As shown in Figure 6B, there is no overlap between HOPX and E cadherin. Given localization outside of epithelial regions, we tested if there was overlap with stromal tissue. To do so, we co-stained with CD73, a known mesenchymal stem cell (MSC) marker, as MSCs play an important role in the generation of cancer-associated fibroblasts and stroma^75^. Combining these results with corresponding tumor sections with H-E staining indicate that HOPX and CD73 are uniquely localized to the tumor stroma. Representative images depicting HOPX, CD73 and HOPX, E cadherin staining for additional samples are shown in Figures S4A and S4B.

It is not surprising that HOPX potentially marks MSCs. Several recent studies have shown HOPX to be associated with other stem cell populations and to play a role in their hierarchy and more importantly maintenance of a stem-cell like state through integration of canonical Wnt and Bmp signaling^70,76,77^. Nonetheless, these results indicate HOPX as a putative novel marker for tumor-associated MSCs. In the patients with poor tumor resectability and prognosis, CD73 and HOPX expression is riddled throughout the tumor tissue (Figure 6B). A typical patient with optimal resectability and low HOPX expression is shown on the left in Figure 6C, whereas a patient with low resectability and high HOPX expression is shown on the right. As can be seen, the tumors with strong evidence of HOPX have very distinct histopathology from those without. This aggressive stromal tumor phenotype provides evidence that patients with poorly resectable tumors have higher levels of stroma that cannot be disentangled from the tumor tissue itself. This suggests one histopathological mechanism for why some tumors are harder to remove from the surrounding stromal tissue. Additionally, the HOPX-CD73 staining indicates that the presence of tumor-associated MSC populations are highly informative of the development of an aggressive stromal phenotype.

## DISCUSSION

We propose the INSPIRE (INferring Shared modules from multiPle gene expREssion datasets) framework for learning a low-dimensional representation (LDR) of multiple gene expression datasets. INSPIRE infers a conserved set of modules and their dependencies across multiple molecular datasets (e.g., gene expression datasets) that contain different sets of genes with a small overlap. We show that INSPIRE outperforms alternative approaches in both synthetically generated datasets and gene expression datasets from ovarian cancer patients. When we applied INSPIRE to nine expression datasets from ovarian cancer studies, which comprises 1,498 patient samples, we identified the stroma/mesenchymal module highly associated with percent stroma and patient survival in the TCGA samples. Our follow-up analysis on this module identifies the *HOPX* gene, which we experimentally validated to be expressed in mesenchymal stem cells (MSCs). *HOPX* is an unusual HOX protein that does not contain a DNA binding domain, and has been implicated in multiple aspects of cardiac and skeletal muscle development through recruitment of histone deacetylases^64–66^. *HOPX* has recently emerged as a marker of numerous stem cell types^70,76,77^. Our results indicate that MSCs are yet another stem cell population marked by *HOPX*. It has been shown that in response to inflammatory cytokines, MSCs release a myriad of growth factors including *FGF*, *EGF*, *PDGF*, and *VEGF*, which promote fibroblasts and endothelial cell differentiation and growth^78^. The tumor MSCs are known contributors to tumor-associated stroma via differentiation to cancer-associated fibroblasts (CAFs)^75^, and may also promote metastasis^79^. *HOPX* could play an important role in this process by acting as a driver given that expression data from *HOPX* knockout mice reveals that many genes in the tumor-associated stroma/mesenchymal module are downstream of *HOPX*. Given the importance of *HOPX* in maintaining a stem cell like state^70^, it is suggestive that *HOPX* expression in the cancer-associated stroma may be maintaining the cancer-associated stroma niche, and could be an attractive target for further functional validation and therapeutic intervention – e.g. if loss of *HOPX* expression in the tumor stroma leads to differentiation of the cancer-associated mesenchymal stem cells.

INSPIRE is a general computational framework, and can be applied to various diseases and different types of molecular data. For example, such as we used applied to integrate mRNA expression datasets from different studies, we can apply it to integrate proteomic data from multiple studies. A future work is to extend INSPIRE such that it can integrate different types of molecular data such as transcriptomic, proteomic, epigenomic and metabolomics data in the same model. In this manuscript, we are applying INSPIRE to integrate microarray data. Since RNA-Seq has been emerging as an important platform for gene expression data profiling, one may want to combine microarray data and RNA-Seq data using INSPIRE. We recommend applying the *voom* normalization method^80^ to read counts when RNA-Seq data are used as input. The *voom* method estimates the mean-variance relationship of the log-counts, generates a precision weight for each observation and enters these into the *limma (Linear Models for Microarray and RNA-Seq Data)* empirical Bayes analysis pipeline. This makes the distributions of the read count data more like a normal distribution, and will make it possible to combine array data with RNA-Seq data using INSPIRE. The authors have shown that the *voom* normalization method has improved statistical properties when applying correlation or linear modeling, which are assumptions in most of the methods being applied to the processed microarray data^80^.

INSPIRE provides a great, effective starting point to learn complex dependencies between genes, because we can learn a gene-level conditional dependence network by using for example the *graphical lasso*^24^ algorithm within each module. There are several other potential next steps to make technical improvement on the proposed INSPIRE framework. One of those is to extend INSPIRE to the case where the latent network is not perfectly conserved across the datasets. We could allow for structured differences characterized by a small subset of modules while we encourage the latent network estimates to be quite similar to each other across datasets. This could be appropriate in many problems where different datasets involve biologically meaningful differences. Another technical improvement is to extend INSPIRE to the setting in which there are no overlapping genes across datasets. For example, one dataset measures the mRNA expression levels of genes and the other dataset measures the protein levels. In this case, we will need to develop a novel method for discovering the correspondences between variables/modules across datasets. Finally, we could exploit the INSPIRE module network information inferred by INSPIRE for imputing the missing variable values in the datasets.

## CONCLUSIONS

In this work, we demonstrate thorough multiple analyses that modules identified by INSPIRE are more biologically coherent across a wide battery of tests of biological significance, including MSigDB pathway enrichment, ChEA transcription factor regulatory networks, and enrichment for known OV CNV tumor drivers. Importantly, the INSPIRE latent variables can be used to predict disease phenotypes or clinical outcome, identify patient subtypes, and when integrated with multiple data modalities, resolve the importance of a specific gene expression module for understanding the mesenchymal subtype in ovarian cancer. Furthermore, when integrated with functional studies of *HOPX* in mice along with immunohistochemistry on multiple patient samples, our analysis suggests an important role for the *HOPX* associated module in maintaining a population of tumor associated mesenchymal stem cells in patients with aggressive stromal components to their tumors.

The effective joint learning strategy of the INSPIRE algorithm makes it possible to integrate datasets containing different sets of genes into a single network framework, which was impossible in the existing network inference approaches. This component of INSPIRE should greatly increase the applicability of LDR learning algorithms to genomics problems where sample size provided by a single dataset is not large enough to learn a robust set of modules and module dependencies. In addition, inferring a network structure among pathways from high-dimensional molecular data is an important and open problem in biology, but is hampered by the need for very large sample sizes. INSPIRE would increase the applicability of network analysis by leveraging existing data, and eliminate the cost of regenerating data from the same samples using different platforms.

## METHODS

### Expression data preprocessing

We downloaded the gene level processed expression data (level 3) for TCGA ovarian cancer from the Firehose pipeline as of the March, 2014 analysis freeze^81^ for all three platforms available for ovarian cancer (Affymetrix U133A, Agilent g4502, Human Exon array). We first removed potential plate level batch effects with ComBat^82^ for all expression datasets. As was done in the TCGA ovarian cancer study^13^, we combined the three separate expression measurements for each of 11864 genes to produce a single estimate of gene expression level by performing a factor analysis across the three studies. All data are log transformed. For other datasets, we downloaded the raw cell intensity files (CEL) for Affymetrix U133 Plus 2.0 and U133A arrays (Affymetrix, Santa Clara, CA, USA) from the Gene Expression Omnibus^34^ for accessions: GSE14764^83^, GSE26712^84^, GSE6008^85^, GSE18520^86^, GSE19829^29^, GSE20565^87^, GSE30161^88^, GSE9899^45^. Expression data were then processed using MAS5.0 normalization with the ‘Affy’ Bioconductor package^89^ and mapped to Entrez gene annotations^90^ using custom chip definition files (CDF)^91^ which was followed by natural log transformation of MAS5.0 normalized intensities. The expression data was then Z-transformed so that each gene has zero mean and unit variance across the samples within each dataset. As stated in Tibshirani (1996)^92^, Z-transformation of expression data is a standard practice for any method that uses a sparsity tuning parameter so that the sparsity tuning parameter is invariant to the scale of the variables, particularly before applying a penalized regression technique such as lasso (*L*_1_ penalty) or ridge (*L*_2_ penalty)^92–96^. Since the graphical model likelihood is indeed equivalent to multiple coupled regression likelihoods, this is generalized to the network estimation problem where we optimize a graphical model likelihood^11,36,97–102^.

### Copy number variation (CNV) data processing

We downloaded the CNV data from 488 ovarian cancer patients in the TCGA cohort from the cBio Cancer Genomics Portal web page^103^. We used R package *cgdsr* to download the data. The 16,597 CNV levels in the downloaded data were derived from the copy-number analysis algorithm GISTIC^104^, and indicate the copy-number level per gene. CNV level ‘-2’ is a deep loss, possibly a homozygous deletion, ’‐1 ’ is a shallow loss (possibly heterozygous deletion), ‘0’ is diploid, ‘1’ indicates a low-level gain, and ‘2’ is a high-level amplification.

### INSPIRE learning algorithm

We present the INSPIRE (<u>**IN**</u>ferring <u>**S**</u>hared modules from multi<u>**p**</u>le gene exp<u>**RE**</u>ssion datasets) method to extract a compact description of high-dimensional gene expression data by learning a set of *k modules and their dependencies* from *Q* gene expression datasets. The technical novelty of the INSPIRE is that it provides a flexible model that does not require the *Q* datasets to have exactly the same set of genes (e.g., different microarray platforms). INSPIRE takes *Q* expression matrices as input and learns how genes are assigned to modules, the latent (unobserved) variables each representing a module, and the dependencies among the latent variables, through an iterative procedure described in detail below. Each latent variable represents the activity level of a certain biological process or a regulatory module. In the sections that describe the probabilistic model and the learning algorithm, we will refer them to as ‘latent variables’ because that is a commonly used term to refer to hidden, unobserved variables in the statistical domain. Inferring the latent variables by using the INSPIRE method is an effective way to obtain a *low-dimensional features* for prediction tasks (e.g., predicting histopathological phenotypes) or clustering (e.g., patient stratification) (Figure 1).

INSPIRE uses a formal probabilistic graphical model, specifically the Gaussian graphical model (GGM), to model the relationships between genes and latent variables, and the conditional dependence relationships among the latent variables. A GGM is a popular probabilistic graphical model for representing the *conditional dependency network* among a set of continuous-valued random variables. In a GGM, the variables connected by an edge are *conditionally dependent* to each other given all the other variables in the model^105,106^. For example, in a simple latent network shown in Figure 1, five *latent variables* (*L*_1_,…, *L*_5_) have mutual dependencies. So, let L = {*L*_1_,…,*L*_5_} ~*N*(0, ∑_L_), then non-zero pattern of ∑_L_^‒1^ corresponds to the conditional dependencies among the latent variables, namely the topology of the network. That means, since *L*_1_ and *L*_2_ are connected to each other for example, knowing’s expression level gives information about *L*_2_’s expression level, even when we know the expression levels of all the other latent variables, which indicates a *direct* dependency between *L*_1_ and *L*_2_. We refer to the observed variables that stem from the same latent variable as a *module*. As an example, genes *G*_1_,*G*_2_,*G*_3_ in Figure 1 form a module since they are associated with the same latent variable *L*_1_. Below, we provide a mathematical formulation of the INSPIRE probabilistic model and the learning algorithm.

Let *X*^1^,…,*X*^Q^ be a set of *Q* expression datasets where the *q*th dataset 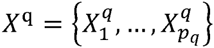 contains the expression levels of *p*_*q*_ genes across *n*_*q*_ samples and each of 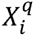 is a row vector of size*n*_*q*_. Let *L*^1^,…,*L*_Q_ be a set of matrices where each *L*^q^ is associated with a dataset and consists of *k* latent variables.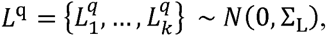, where ∑_L_ is a *K* × *K* covariance matrix. These latent variables can be viewed as a lower-dimensional representation (LDR) of expression data and ∑_L_ represents the dependencies among the features. We assume that ∑_L_ is conserved across the *Q* datasets. Each gene is associated with exactly one of the *k* latent variables as represented by the directed edge between a gene and a latent variable in Figure 1. The total number of unique genes across all *Q* datasets is *p*_*T*_; and each data matrix *X*^*q*^ contains samples from a different subset of *p*_*q*_ genes (*p*_*q*_ ≤ *p*_*T*_). Let Z be a *p*_*T*_ × *k* matrix indicating which of the *k* modules each of *p*_*T*_ genes belongs to, such that ∀*i*,*j Z*_*ij*_ ∈ {0,1} and 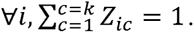.Each observed dataset *X*^*q*^ is generated by the multivariate Gaussian distribution *X*^q^ | Z^q^L^*q*^, *σ*^2^ ~ *N*(Z^*q*^*L*^*q*^, *σ*^2^), where *Z*^*q*^ is a *p*_*q*_ × *K* matrix composed of the rows of Z corresponding to the *p*_*q*_ genes contained by the dataset *X*_*q*_.Here, we refer to a set of genes that correspond to the same latent variable as a *module* where determines the module tightness. As an example, the *j* th module *M*_*j*_ can be defined as 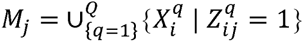. Thus, *Z* defines the module assignment of all unique genes in all *Q* datasets into *k* modules. Each gene belongs to exactly one module. We choose hard assignment of genes to modules (∀*i*, ∃!*c* : *Z*_*ic*_ = 1) to reduce the number of parameters. Soft assignment is a straightforward extension where we relax the constraint ∀*i*,*j Z*_*ij*_ ∈ {0,1} to ∀*i*,*j* 0 ≤ *Z*_*ij*_ ≤ 1.

INSPIRE jointly learns the latent variables *L* = [*L*^1^,…,*L*^Q^] eachcorrespondingtoa module; the module assignment indicator *Z*; and the feature dependence network ∑_*L*_^‒1^. Given *Q* datasets *X*^1^,…,*X*^*Q*^, where *X*^*q*^( ∈ ℝ^{*p*_*q*_×*n*_*q*_}^)contains *n*_*q*_ observations on *p*_*q*_ genes and 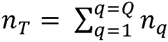, INSPIREaimstolearn the following:

‐ *L*^*q*^ ∈ ℝ^{*K* × *n*_*q*_}^ for each *q*(∈ {1,…,*Q*}) containing the values on *k* features in *n*_*q*_ samples in *X*^*q*^
‐ Z | ∑*Z*_*i*_ = 1, a binary vector for each *i*(∈ {1,…,*p*_*T*_}) specifying the module membership of the *i*th gene in one of the *k* modules; and
‐ *Θ*_*L*_ ( ∈ ℝ^{*k* × *k*}^) denoting the estimate of the inverse covariance matrix of the features, i.e. ∑_*L*_^‒1^.

We address our learning problem by finding the joint maximum a posteriori (MAP) assignment to all of the optimization variables – *L*, *Z*, and *Θ*_*L*_. This means that we optimize the joint log-likelihood function of the *Q* data matrices, with respect to *L*,*Z*, and *Θ*_L_( ≻ 0).Given the statistical independence assumption that genes in a dataset *X*^*q*^ are statistically independent to one another given the latent variables *L*^*q*^, the joint log likelihood can be decomposed as follows:

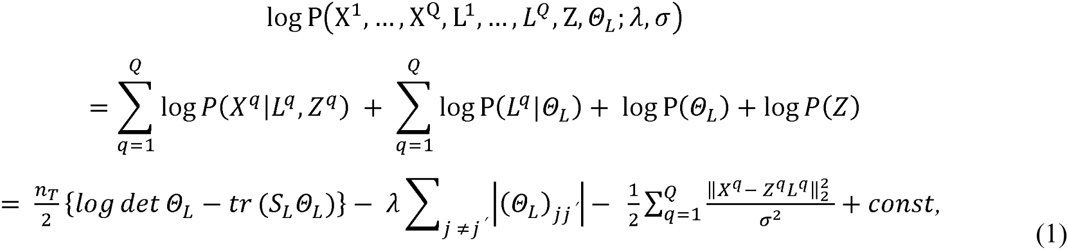

where 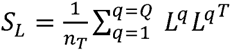 is the empirical estimate of the covariance matrix ∑_*L*_ and *λ* is a positive tuning parameter that adjusts the sparsity of *Θ*_*L*_. We assume a uniform prior distribution over *Z*, which makes log *P*(*Z*) constant.

We use a *coordinate ascent procedure* over three sets of optimization variables –,*Z*, and *Θ*_*L*_.We iteratively estimate each of the optimization variables until convergence.

<u>**Learning L**</u> To estimate *L*^1^,…, *L*^*Q*^ from equation 1 given *Z* and **Θ**_*L*_, we solve the following problem:

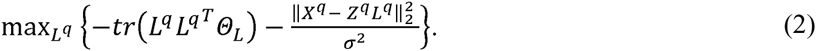

Setting the derivative of the objective function in equation 2 to zero with respect to *L*^*q*^ leads to:

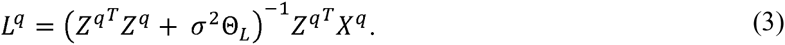

<u>**Learning Z**</u> In order to estimate *Z* given *L*^1^,…, *L*^Q^, we solve the following optimization problem:

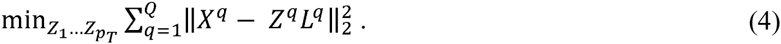

In the hard assignment paradigm that we follow throughout this paper, equation 4 assigns gene *P*_*i*_ to module *c* ∈ {1,…, *K*} that minimizes the Euclidean distance computed using all samples from the datasets containing the gene *P*_*i*_.

<u>Learning *Θ*,_*L*_</u> To estimate *Θ*_*L*_ given *L*^*1*^,…,*L*^*Q*^, we solve the following optimization problem:

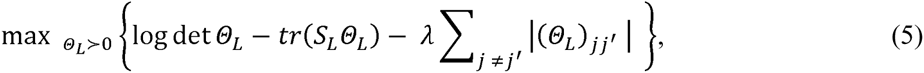

where the constraint *Θ*_*L*_ *≻* 0 restricts the solution to the space of positive definite matrices of size *k* × *k*, and 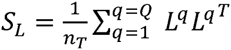 is the empiricalcovariance matrixof *L*. Based on the estimated value of *L*, equation 5 can be solved by the graphical lasso^24^, a well-known algorithm for learning the structure of a Gaussian graphical model (GGM).

We iteratively estimate each of the optimization variables until convergence. Since our objective is continuous on a compact level set, based on Theorem 4.1 in Tseng (2001)^107^, the solution sequence is defined and bounded. Every coordinate group reached by the iterations is a stationary point of INSPIRE objective function. We also observed that the value of the objective likelihood function monotonically increases.

### Data imputation

To our knowledge, there are no published methods for learning modules and their dependencies from multiple datasets that contain different sets of genes (Figure 1). Thus, we adapted the state-of-the-art methods (which can run on a single dataset) by imputing the missing values on genes that are not presented in each of the datasets, and applied these methods to the imputed data. These are the ‘Imp‐‐’ methods in Table 1. We employed the iterative PCA algorithm to generate the imputed data for all ‘Imp‐‐’ methods and initializing INSPIRE. The results were robust to the imputation method; INSPIRE method consistently outperformed alternative approaches when other imputation methods were used. We used CRAN R package missMDA^108^ to generate the imputed data.

### Initialization of the INSPIRE latent variables

INSPIRE is an iterative learning algorithm that consists of three update steps, Equations (3)-(5), to learn the following sets of parameters: ***L*** – values on the latent variables, ***Z*** – gene-module assignments, and *θ*_*L*_–the dependency network among the latent variables. So we need to have some starting point, i.e., initial values on any of these three sets of parameters. SLFA and MGL are also iterative learning algorithms that require a starting point. Therefore, for INSPIRE, SLFA and MGL, we used the same initial gene-module assignments obtained by running the *k*-means clustering algorithm on the imputed data (see above) because the imputed data contain all genes and all samples.

To be more specific, the authors of the MGL algorithm suggested to initialize MGL with *k*-means centroids, and we followed that approach for the MGL variants (MGL1, ImpMGL, and InterMGL) in our experiments. Given that INSPIRE is an extension to MGL for multi-data setting, to directly test whether the INSPIRE outperforms MGL, we used the output of MGL as a starting point for INSPIRE. The authors of the SLFA algorithm did not specify any initialization method; so for a fair comparison among all these methods, we used the same initial gene-module assignments for SLFA and MGL - the centroids obtained by running the *k*-means clustering algorithm on the imputed data. The result of the *k*-means clustering algorithm also depends on the initial clusters which are randomly determined. So, to rule out the possibility to make a conclusion based on a particular set of initial parameters, for every experiment on comparison across methods, we performed 10 runs with different initial parameters (i.e., different random initial clusters in the *k*-means clustering algorithm) and presented the average results.

### Runtime of INSPIRE on gene expression datasets

Running INSPIRE with the module count parameter *k* = 90 and the sparsity tuning parameter *λ* = 0.1 in our application on nine datasets (Table S2) with a total number of *p* ≅ 20,000 genes and *n* ≅ 1,500 samples took 13.7 minutes on a machine with an Intel(R) Xeon(R) E5645 2.40GHz CPU and 24GB RAM, once the latent variables are initialized. As mentioned above, for initialization of the latent variables, we used the module graphical lasso (MGL)^11^ method on the imputed data, which took 10.2 minutes on the same machine.

### Synthetic data generation

We synthetically generated data based on the joint distribution in equation 1. We first generated the sparse *k* × *k* inverse covariance matrix ∑_*L*_^‒1^ by creating a *k* × *k* matrix G as

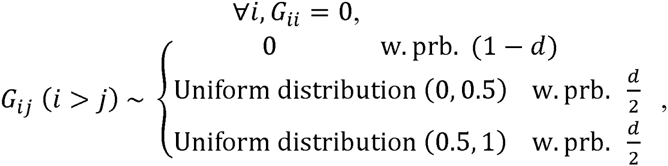

and letting ∑_*L*_^‒1^ = *G* + *G*^*T*^ = *G* + *G*^*T*^ so that ∑_*L*_^‒1^ is symmetric. We set ∀*i*, *G*_*ii*_ = *ϵ* afterwards by selecting *ϵ* such that the resulting matrix ∑_*L*_^‒1^ is positive definite. *d ∈* [0,1] controls the density of ∑_*L*_^‒1^ and the results we reported from synthetic data experiments were generated using *k* = 10 and *d* = 0.2. The results were consistent for varying values of *k* and *d*.

Then, we generated the latent variables *L* = {*L*_1_,…, *L*_*k*_} from *L* ~ N(0,∑_*L*_) and we randomly generated a binary *P*_*T*_ × *k* matrix *Z* of module assignments which randomly assigns each of *P*_*T*_ genes to exactly one of the latent variables. Then we generated a high-dimensional data matrix *X* of *p*_*T*_ genes from the distribution *X* | *ZL*,*σ*^2^ ~ *N*(*ZL*,*σ*^2^) and selected a portion of the samples and genes in *X* to form a smaller dataset that we call ‘Dataset1’. Then we selected the remaining samples and a portion of the genes from *X* to form a second ‘Dataset2’.

We considered three simulated settings that correspond to different amount of overlapping genes (Figure S1A). Each setting is characterized by [*0L*,*D1*,*D2*] where *OL* denotes the number of genes that are present in both Dataset1 and Dataset2, is the number of genes that are present only in Dataset1, and *D2* means the number of genes that are present only in Dataset2. The settings we consider are, and,where the sample sizes of Dataset1 and Dataset2 are 20 and 30,respectively (Figure S1A). [250,0,0] means that all genes are shared between the two datasets. We repeated the generation of data *X* 20 times in each of the three settings, and presented the mean of the results for each method in (Figure 3A-C). We show the *p*-values on the bars that represent the statistical significance of the difference between each method and INSPIRE across 20 different data instantiations.

Figure S1A illustrates the two datasets in each of these three settings. In each rectangle, each row represents a variable, and each column represents a sample. For simplicity in presentation of the evaluation results, we set *D*2 = 0. The results were consistent for varying *D*2. We note that *D*2 ≅ 0 assumption holds in many real-world settings we are interested in, where the newer technology contains almost all of the genes in the older technology. We demonstrate this real-world situation in the second set of experiments on the ovarian cancer expression data (Figure 4B).

### Comparison of the scalability across all six methods in simulation experiment

We precisely measured the runtimes of six methods – GLasso, UGL, SLFA, WGCNA, MGL (Table 1) and INSPIRE – when running on the synthetic data with varying numbers of genes (*p*); *p* = 300,*p* = 1,500,*p* = 3,000.We generated the data exactly the same way as in the simulation experiments. We used 50 as sample size (20 samples in Dataset1 and 30 samples in Dataset2). We tested these methods on the ‘Imp‐‐‐’ setting where we imputed the missing data before applying the algorithms, because 5 of these methods (except INSPIRE) cannot accommodate multiple datasets. We used varying sparsity tuning parameters in the interval of, exactly the same set of values that we used for choosing λ(via cross-validation tests) in our experiments. The runtimes of these methods are known to grow cubically or at least quadratically depending on the availability of a special efficient technique for the method^109^ with increasing *p* (when gene-level dependencies are learned – GLasso and UGL), or *k* (when module level dependencies are learned – SLFA and MGL). Also, WGCNA grows quadratically with increasing *p* since it includes correlation computation and hierarchical clustering. Therefore, we determined that the methods whose runtime is >10 hours for *p* = 3,000 are not scalable enough to be useful on genome-wide analysis. Since the runtimes of the methods except MGL, WGCNA and INSPIRE already exceeded 10 hours at *p* = 3,000 (Figure S9A), it is clear that all methods other than MGL, WGCNA and INSPIRE are too slow to be used when *p* > 3,000 and >500 hours when *p* is near 20,000 (see the trend line in Figure S9B). We note that we increased the module count (*k*) with increasing *p* such that the average number of genes in a module is always 30, and SLFA was unable to run for *p* > 1,500 where the module count (*k*) exceeded the sample size (50). Figure S9A-B indicate that GLasso, UGL and SLFA are not practically useful to be used on genome-wide expression datasets, and furthermore, they do not perform well on smaller synthetic data on which we ran all six methods (Figure 3). Thus, we excluded GLasso, UGL and SLFA for the evaluation on the genome-wide expression datasets. The runtime measurements were done on a very powerful machine with an Intel(R) Xeon(R) E7-8850 v2 @ 2.30GHz CPU and 528 GB RAM.

### Computing the cross validation test log-likelihood

We performed a 5-fold CV to choose *λ* for INSPIRE and each of the competing methods in our experiments to evaluate INSPIRE (synthetic data experiments and the experiments with two gene expression datasets). We measured the CV test log-likelihood on the test data portion of the first dataset (Dataset1 or OV1 which contains all or almost all genes) in each fold, which was common test data across all methods. For each of the 5 test folds, we computed the test data log-likelihood of the *p* × *p* gene-level dependency matrix that is computed using the dependencies among the latent variables (representing modules) inferred by each of the INSPIRE and its competitors, where *p* is the total number of genes in the two datasets. For the methods that optimize a non-convex objective function, we averaged the CV test log-likelihoods across multiple runs with different initial assignment of genes to modules. We tested a range of sparsity tuning parameter values (*λ*), and observed the “cup-shaped” underfitting/overfitting pattern in the *λ* (x-axis) vs. average CV test log-likelihood (y-axis) curves for all methods, as expected.

### Evaluation of learned network in synthetic data experiments

In the synthetic data experiments, the correspondence between the modules in a learned model and the modules in the true model is not clear because each method can end up having different optimal number of modules, even if they started with the same number of initial modules. Therefore, we compared the methods in terms of the accuracy of the *p* × *p* gene-level dependency matrix that is computed using the dependencies among the modules inferred by each of the INSPIRE and its competitors, where *p* is the total number of genes in the two datasets.

### Measuring the significance of difference between INSPIRE and 13 competing methods

We repeated the synthetic data generation 20 times in each of the three settings, and presented the average results with the Wilcoxon signed rank test *p*-value measuring the significance of differences based on the Wilcoxon signed rank test. More specifically, it measures the probability that the corresponding method gave a better result in terms of mean rank than INSPIRE across 20 different data instantiations.

### Comparison of the prediction performance with alternative methods

We compared INSPIRE with principal component analysis (PCA) and subnetwork analysis method^15^ based on how well each method can predict each of the six phenotypes (resectability as defined by 0 cm of residual tumor vs. >0 cm of residual tumor after surgery, survival time, and four manually curated histologic phenotypes) from The Cancer Genome Atlas (TCGA) data. We used the *lasso*^110^ (L_1_ regularized linear regression) for predicting the continuous-valued phenotype (percent stroma), L_1_ regularized logistic regression for predicting binary phenotypes (stroma type, vessel formation, invasion pattern, and residual tumor), and L_1_ regularized Cox regression for predicting survival. The prediction performance was measured in left-out data via leave-one-out cross validation (LOOCV) tests for histologic phenotypes that have relatively less number of samples (~100), and 50-fold cross-validation for resectability and survival that have larger number of samples (~500). The sparsity tuning parameter *λ* was chosen within training data by performing LOOCV tests, which is a standard way of choosing *λ*^110^. For a fair comparison with PCA^20^ and the subnetwork method^15^, we used top 90 principal components, and 90 subnetworks that are most correlated with the phenotype, respectively. The subnetwork analysis method runs on binary phenotypes, but ‘percent stroma’ is continuous-valued; so, to make the subnetwork method work on this phenotype, we binarized the values by making >50% to be 1 and >50% be 0.

### Learning subtypes based on the INSPIRE latent variables

We used the *k*-means clustering algorithm on the INSPIRE latent variables, each of which corresponds to a module, to cluster patients into four subtypes. We chose four as the number of subtypes to make it comparable to alternative subtyping methods (TCGA study^13^ and the NBS method^35^). Since *k*-means is non-deterministic, the resulting subtypes could depend on the starting point of the subtype assignments. In order to get the most coherent groups of patients, we ran *k*-means 10 times with different random initial assignments of the patients into subtypes, and chose the clustering which gives the lowest within cluster sum of squares.

### Supervised model to predict tumor resectability

We trained supervised models of tumor resectability using different combinations of the POSTN expression and the latent variables corresponding to module 5 and module 6 in TCGA ovarian cancer data for 489 patients to predict 0 cm of residual tumor vs. >0 cm of residual tumor. The proportion of the sub-optimally debulked patients was 62% (= 139/223) in Tothill^45^ and was 77% (= 378/489) in TCGA^13^. Logistic regression was used to train the models. Five distinct models were constructed: 1) a model with only the POSTN expression, 2) a model with only the latent variable corresponding to module 5, 3) a model with only the latent variable corresponding to module 6, 4) a model with POSTN expression and the latent variable corresponding to module 5, and 5) a model with the latent variables corresponding to module 5 and module 6. We trained each of those models along with (Figure 7) and without (Figure S5) the clinical covariates of age and stage. Performance was determined based on the results of each fitted model in the Tothill^45^ data in terms of the Area Under the Curve (AUC) measure from a Receiver Operator Characteristic (ROC) curve (Figure 7 and Figure S5).

### Extraction of tumor histologic phenotypes from TCGA images

We manually curated multiple tumor histopathology features from image data on H-E staining of ovarian tumor section from TCGA. We primarily used 98 randomly sampled patients to test the association between tumor histopathology features and the latent variables learned by INSPIRE. Features were curated in a blinded fashion. Five histopathological features were evaluated including percent stroma, percent tumor, vessel formation, stroma type, and pattern of invasion. Percent tumor was defined as the percent area involved by viable neoplastic cells across the entire slide while percent stroma was the percent area of fibrous tissue (fibroblasts and collagen). Vessel formation was scored as minimal, moderate, or abundant based on the number of formed vessels identified at 100X magnification. Stroma type was defined as fibrous (dense collagen with relatively fewer fibroblasts) or desmoplastic (many fibroblasts embedded in a loose, myeloid extracellular matrix). Pattern of invasion related to how the neoplastic cells interacted with the surrounding stroma and was scored as expansile, infiltrative, papillary, or mixed. Expansile invasion was characterized by cohesive tumor cells growing in a cluster with relatively well-circumscribed borders with the surrounding stroma while infiltrative invasion included tumor cells which grew in small nests or tentacles with abundant stroma surrounding the individual tumor cells. Tumors classified as having papillary invasion had abundant fibro-vascular cores upon which the neoplastic cells grew in arborizing branches. Mixed invasion patterns were identified and classified as such.

### Immunohistochemistry

Ten patients were sampled for staining based on either having good tumor resection and survival (> 3 year survival, optimal debulking with residual tumor < 1cm) vs. poor tumor resection and survival (< 3 years survival, > 1 cm residual tumor). Tissue and clinical information were collected with patient consent by the University of Washington Gynecologic Oncology Tissue Bank under approval from the human subjects division (IRB 27077). Tumor tissue was collected at the time of primary surgery and flash frozen in liquid nitrogen, transported to the lab and stored at ‐80 C. The 17 frozen block was cryo-sectioned and one 8 mm section placed on a charged slide for IHC testing and H-E staining.

Frozen tissue slices fixed to glass slides were allowed to thaw at room temp for ten min. Slides were fixed in a Coplin jar in cold acetone for ten min at ‐20C. Slides were removed from acetone and placed tissue side up on a shaker. PBS was added to the slide (1mL, enough to cover tissue slice) for five min shaking. PBS wash was repeated for a total of two five min washes. After final wash, PBS was poured off the slide and tissues were blocked with 2% milk/PBS (Carnation Instant Nonfat Dry Milk dissolved in PBS) for one hour at room temperature, while shaking. Blocking solution was removed and primary antibody added, diluted in 1% milk. Antibody dilutions were per manufacturers recommendations. Slides were allowed to incubate overnight at 4°C while shaking with primary antibody. If primary antibody was conjugated to fluorescent molecule, slides were also incubated in the dark overnight. Slides were washed three times with PBS at room temperature. Secondary antibody was diluted in 1% milk/PBS and incubated at room temperature for 30 min, shaking. Slides were then washed with PBS for 10 min, three times. Nuclear stain diluted in PBS was added to tissues. Either Dapi (300ng/mL, Sigma-Aldrich, catalog #D9542) or Sytox Green Nuclear Stain (Life Technologies, catalog # S7020) was used depending on the secondary antibodies used for staining. Last PBS wash was done at room temperature for 5min. Coverslips were mounted to slides using Fluoroshield (Sigma-Aldrich, catalog # F6182) and sealed with clear nail polish. Images were taken on a Nikon TiE Inverted Widefield Fluorescence High Resolution Microscope.

Primary Antibodies used: Anti-E Cadherin antibody conjugated to Allophycocyanin (Abcam, catalog # ab99885), Hop Antibody (Santa Cruz, catalog # sc-30216), Anti-CD73 antibody (Abcam, catalog # ab54217), GCS-a-1 Antibody (Santa Cruz, sc-23801)

Secondary Antibodies used: CD73 antibody was detected with Goat anti-mouse IgG-FITC (Santa Cruz, catalog # sc-2010). When co-stained with CD73, HOPX was detected with Donkey anti-rabbit IgG-CFL 647 (Santa Cruz, catalog # sc-362291). When co-stained with E Cadherin, HOPX antibody was detected with Chicken anti-rabbit IgG H&L FITC (Abcam, catalog # ab6825).

### Analysis of immunohistochemistry

Fluorescence images were analyzed using ImageJ^111^ and the plugin JACoP was used for co-localization analysis.

## DECLARATIONS

### List of abbreviations

INSPIRE: INferring Shared modules from multiPle gene expREssion datasets
TCGA: The Cancer Genome Atlas
LDR: Low-Dimensional Representation
WGCNA: Weighted Gene Co-expression Network Analysis
GGM: Gaussian Graphical Model
EMT: Epithelial-Mesenchymal Transition
BIC: Bayesian Information Criterion
PPI: Protein-Protein Interaction
MSigDB: Molecular Signatures DataBase
PCA: Principal Component Analysis
PC: Principal Component
TOM: Topological Overlap Measure
MGL: Module Graphical Lasso
SLFA: Structured Latent Factor Analysis
GLasso: Graphical Lasso
UGL: Unknown Group L1 regularization
CV: Cross-Validation
TF: Transcription Factor
ChEA: ChIP Enrichment Analysis
GEO: Gene Expression Omnibus
CNV: Copy Number Variation
NBS: Network-Based Stratification
SAM: Significance Analysis of Microarrays
OV: OVarian cancer
AUC: Area Under the Curve
MSC: Mesenchymal Stem Cell
CAF: Cancer-Associated Fibroblasts
MAP: Maximum A Posteriori
LOOCV: Leave-One-Out Cross Validation
ROC: Receiver Operator Characteristic
GISTIC: Genomic Identification of Significant Targets in Cancer

### Ethics approval and consent to participate

All input data of INSPIRE are publicly available through the Gene Expression Omnibus (GEO) web page^112^ and The Cancer Genome Atlas (TCGA). For experimental validation on HOPX, we used frozen tissue slices fixed to glass slides from ten patients with ovarian cancer for immunohistochemistry staining. The tumor tissue and associated clinical variables were obtained from an institutional tumor bank, which prospectively collected specimens and clinical information for subjects who provided informed consent under an IRB-approved protocol (University of Washington IRB 27077).

### Consent for publication

Not applicable.

### Availability of data and materials

INSPIRE is freely available as an R package in the CRAN repository. The processed expression data used in the study, the inferred INSPIRE models, histopathologic features of 100 TCGA H&E stained images, and the results of our immunohistochemistry staining experiments are available on our website^14^.

### Author’s contributions

SC, BAL and SIL designed the statistical methods and various analyses. SC developed the algorithm, and performed computational experiments. SC, BAL, SB, MR, RDH and SIL wrote the manuscript. RDH and SB designed and performed the immunohistochemistry experiment. MR extracted histopathologic features from the 100 H&E stained ovarian tumor section images obtained through TCGA. CD interpreted the results on the molecular basis for resectability. All authors read and approved the manuscript.

## REFERENCES

Unsupervised Feature Learning and Deep Learning Tutorial.

Langkvist, M., Karlsson, L. & Loutfi, A. A review of unsupervised feature learning and deep learning for time-series modeling. Pattern Recognit. Lett:. 42, 11–24 (2014).

LeCun, Y., Bengio, Y. & Hinton, G. Deep learning. Nature 521, 436–444 (2015).

Szegedy, C., Toshev, A. & Erhan, D. Deep Neural Networks for Object Detection. in Adv. Neural Inf. Process. Syst. 2553–2561 (2013).

Lecun, Y., Bottou, L., Bengio, Y. & Haffner, P. Gradient-based learning applied to document recognition. Proc. IEEE 86, 2278–2324 (1998).

Margolin, A. A., Bilal, E., Huang, E., Norman, T. C., Ottestad, L., Mecham, B. H., Sauerwine, B., Kellen, M. R., Mangravite, L. M., Furia, M. D., Vollan, H. K. M., Rueda, O. M., Guinney, J., Deflaux, N. A., Hoff, B., Schildwachter, X., Russnes, H. G., Park, D., Vang, V. O., Pirtle, T., Youseff, L., Citro, C., Curtis, C., Kristensen, V. N., Hellerstein, J., Friend, S. H., Stolovitzky, G., Aparicio, S., Caldas, C. & Børresen-Dale, A.-L. Systematic analysis of challenge-driven improvements in molecular prognostic models for breast cancer. Sci. Transl. Med. 5, 181re1 (2013).

Cheng, W.-Y., Yang, T.-H. O. & Anastassiou, D. Development of a Prognostic Model for Breast Cancer Survival in an Open Challenge Environment. Sci. Transl. Med. 5, 181ra50 (2013).

Langfelder, P. & Horvath, S. WGCNA: an R package for weighted gene co-expression network analysis. BMC Bioinformatics 9, 559 (2008).

Sherlock, G. Analysis of large-scale gene expression data. Brief. Bioinform. 2, 350–62 (2001).

Lee, S.-I. & Batzoglou, S. in Adv. Neural Inf. Process. Syst. 16 (eds. Thrun, S., Saul, L. & Schölkopf, B.) (MIT Press, 2004).

Celik, S., Logsdon, B. A. & Lee, S.-I. Efficient Dimensionality Reduction for High-Dimensional Network Estimation. in ICML (2014).

Koller, D. & Friedman, N. Probabilistic Graphical Models: Principles and Techniques. (The MIT Press, 2009).

Bell, D., Berchuck, A., Birrer, M., Chien, J., Cramer, D. W., Dao, F., Dhir, R., DiSaia, P., Gabra, H., Glenn, P., Godwin, A. K., Gross, J., Hartmann, L., Huang, M., Huntsman, D. G., Iacocca, M., Imielinski, M., Kalloger, S., Karlan, B. Y., Levine, D. A., Mills, G. B., Morrison, C., Mutch, D., Olvera, N., Orsulic, S., Park, K., Petrelli, N., Rabeno, B., Rader, J. S., Sikic, B. I., Smith-McCune, K., Sood, A. K., Bowtell, D., Penny, R., Testa, J. R., Chang, K., Dinh, H. H., Drummond, J. A., Fowler, G., Gunaratne, P., Hawes, A. C., Kovar, C. L., Lewis, L. R., Morgan, M. B., Newsham, I. F., Santibanez, J., Reid, J. G., Trevino, L. R., Wu, Y.-Q., Wang, M., Muzny, D. M., Wheeler, D. A., Gibbs, R. A., Getz, G., Lawrence, M. S., Cibulskis, K., Sivachenko, A. Y., Sougnez, C., Voet, D., Wilkinson, J., Bloom, T., Ardlie, K., Fennell, T., Baldwin, J., Gabriel, S., Lander, E. S., Ding, L., Fulton, R. S., Koboldt, D. C., McLellan, M. D., Wylie, T., Walker, J., O’Laughlin, M., Dooling, D. J., Fulton, L., Abbott, R., Dees, N. D., Zhang, Q., Kandoth, C., Wendl, M., Schierding, W., Shen, D., Harris, C. C., Schmidt, H., Kalicki, J., Delehaunty, K. D., Fronick, C. C., Demeter, R., Cook, L., Wallis, J. W., Lin, L., Magrini, V. J., Hodges, J. S., Eldred, J. M., Smith, S. M., Pohl, C. S., Vandin, F., Raphael, B. J., Weinstock, G. M., Mardis, E. R., Wilson, R. K., Meyerson, M., Winckler, W., Verhaak, R. G. W., Carter, S. L., Mermel, C. H., Saksena, G., Nguyen, H., Onofrio, R. C., Hubbard, D., Gupta, S., Crenshaw, A., Ramos, A. H., Chin, L., Protopopov, A., Zhang, J., Kim, T. M., Perna, I., Xiao, Y., Zhang, H., Ren, G., Sathiamoorthy, N., Park, R. W., Lee, E., Park, P. J., Kucherlapati, R., Absher, D. M., Waite, L., Sherlock, G., Brooks, J. D., Li, J. Z., Xu, J., Myers, R. M., Laird, P. W., Cope, L., Herman, J. G., Shen, H., Weisenberger, D. J., Noushmehr, H., Pan, F., Triche Jr, T., Berman, B. P., Van Den Berg, D. J., Buckley, J., Baylin, S. B., Spellman, P. T., Purdom, E., Neuvial, P., Bengtsson, H., Jakkula, L. R., Durinck, S., Han, J., Dorton, S., Marr, H., Choi, Y. G., Wang, V., Wang, N. J., Ngai, J., Conboy, J. G., Parvin, B., Feiler, H. S., Speed, T. P., Gray, J. W., Socci, N. D., Liang, Y., Taylor, B. S., Schultz, N., Borsu, L., Lash, A. E., Brennan, C., Viale, A., Sander, C., Ladanyi, M., Hoadley, K. A., Meng, S., Du, Y., Shi, Y., Li, L., Turman, Y. J., Zang, D., Helms, E. B., Balu, S., Zhou, X., Wu, J., Topal, M. D., Hayes, D. N., Perou, C. M., Zhang, J., Wu, C. J., Shukla, S., Sivachenko, A., Jing, R., Liu, Y., Noble, M., Carter, H., Kim, D., Karchin, R., Korkola, J. E., Heiser, L. M., Cho, R. J., Hu, Z., Cerami, E., Olshen, A., Reva, B., Antipin, Y., Shen, R., Mankoo, P., Sheridan, R., Ciriello, G., Chang, W. K., Bernanke, J. A., Haussler, D., Benz, C. C., Stuart, J. M., Benz, S. C., Sanborn, J. Z., Vaske, C. J., Zhu, J., Szeto, C., Scott, G. K., Yau, C., Wilkerson, M. D., Zhang, N., Akbani, R., Baggerly, K. A., Yung, W. K., Weinstein, J. N., Shelton, T., Grimm, D., Hatfield, M., Morris, S., Yena, P., Rhodes, P., Sherman, M., Paulauskis, J., Millis, S., Kahn, A., Greene, J. M., Sfeir, R., Jensen, M. A., Chen, J., Whitmore, J., Alonso, S., Jordan, J., Chu, A., Zhang, J., Barker, A., Compton, C.,Eley, G., Ferguson, M., Fielding, P., Gerhard, D. S., Myles, R., Schaefer, C., Mills Shaw, K. R., Vaught, J., Vockley, J. B., Good, P. J., Guyer, M. S., Ozenberger, B., Peterson, J. & Thomson, E. Integrated genomic analyses of ovarian carcinoma. Nature 474, 609–615 (2011).

INSPIRE web page: http://inspire.cs.washington.edu.

Chuang, H.-Y., Lee, E., Liu, Y.-T., Lee, D. & Ideker, T. Network-based classification of breast cancer metastasis. Mol. Syst. Biol. 3, 140 (2007).

Lee, E., Chuang, H. Y., Kim, J. W., Ideker, T. & Lee, D. Inferring pathway activity toward precise disease classification. PLoS Comput. Biol. 4, (2008).

Taylor, I. W., Linding, R., Warde-Farley, D., Liu, Y., Pesquita, C., Faria, D., Bull, S., Pawson, T., Morris, Q. & Wrana, J. L. Dynamic modularity in protein interaction networks predicts breast cancer outcome. Nat. Biotechnol. 27, 199–204 (2009).

Ravasi, T., Suzuki, H., Cannistraci, C. V., Katayama, S., Bajic, V. B., Tan, K., Akalin, A., Schmeier, S., Kanamori-Katayama, M., Bertin, N., Carninci, P., Daub, C. O., Forrest, A. R. R., Gough, J., Grimmond, S., Han, J. H., Hashimoto, T., Hide, W., Hofmann, O., Kawaji, H., Kubosaki, A., Lassmann, T., van Nimwegen, E., Ogawa, C., Teasdale, R. D., Tegn??r, J., Lenhard, B., Teichmann, S. A., Arakawa, T., Ninomiya, N., Murakami, K., Tagami, M., Fukuda, S., Imamura, K., Kai, C., Ishihara, R., Kitazume, Y., Kawai, J., Hume, D.A., Ideker, T. & Hayashizaki, Y. An Atlas of Combinatorial Transcriptional Regulation in Mouse and Man. Cell 140, 744–752 (2010).

Herschkowitz, J. I., Simin, K., Weigman, V. J., Mikaelian, I., Usary, J., Hu, Z., Rasmussen, K. E., Jones, L. P., Assefnia, S., Chandrasekharan, S., Backlund, M. G., Yin, Y., Khramtsov, A. I., Bastein, R., Quackenbush, J., Glazer, R. I., Brown, P. H., Green, J. E., Kopelovich, L., Furth, P. A., Palazzo, J. P., Olopade, O. I., Bernard, P. S., Churchill, G. A., Van Dyke, T. & Perou, C. M. Identification of conserved gene expression features between murine mammary carcinoma models and human breast tumors. Genome Biol. 8, R76 (2007).

Hotelling, H. Analysis of a complex of statistical variables into principal components. J. Educ. Psychol. 24, 417–441 (1933).

Jiang, D., Tang, C. & Zhang, A. Cluster analysis for gene expression data: A survey. IEEE Trans. Knowl. Data Eng. 16, 1370–1386 (2004).

Chandrasekaran, V., Parrilo, P. A. & Willsky, A. S. Latent variable graphical model selection via convex optimization. Ann. Stat. 40, 1935–1967 (2012).

He, Y., Qi, Y., Kavukcuoglu, K. & Park, H. Learning the dependency structure of latent factors. in NIPS (2012).

Friedman, J., Hastie, T. & Tibshirani, R. Sparse inverse covariance estimation with the graphical lasso. Biostatistics 9, 432–441 (2008).

Marlin, B. M. & Murphy, K. Sparse gaussian graphical models with unknown block structure. in ICML (2009).

Duchi, J. & Gould, S. Projected subgradient methods for learning sparse gaussians. Twenty-fourth Conf. 145–152 (2008). at <http://uai.sis.pitt.edu/papers/08/p153-duchi.pdf>

Hastie, T., Tibshirani, R. & Friedman, J. The Elements of Statistical Learning. Book 2nded, (2001).

Rand, W. M. Objective Criteria for the Evaluation of Clustering Methods. J. Am. Stat. Assoc. 66, 846–850 (1971).

Konstantinopoulos, P. A., Spentzos, D., Karlan, B. Y., Taniguchi, T., Fountzilas, E., Francoeur, N., Levine, D.A. & Cannistra, S. A. Gene expression profile of BRCAness that correlates with responsiveness to chemotherapy and with outcome in patients with epithelial ovarian cancer. J. Clin. Oncol. 28, 3555–3561 (2010).

Lachmann, A., Xu, H., Krishnan, J., Berger, S. I., Mazloom, A. R. & Ma’ayan, A. ChEA: transcription factor regulation inferred from integrating genome-wide ChIP-X experiments. Bioinformatics 26, 2438–44 (2010).

Subramanian, A., Tamayo, P., Mootha, V. K., Mukherjee, S., Ebert, B. L., Gillette, M. a, Paulovich, A., Pomeroy, S. L., Golub, T. R., Lander, E. S. & Mesirov, J. P. Gene set enrichment analysis: a knowledge-based approach for interpreting genome-wide expression profiles. Proc. Natl. Acad. Sci. U. S. A. 102, 15545–50 (2005).

Vastrik, I., D’Eustachio, P., Schmidt, E., Joshi-Tope, G., Gopinath, G., Croft, D., de Bono, B., Gillespie, M., Jassal, B., Lewis, S., Matthews, L., Wu, G., Birney, E. & Stein, L. Reactome: a knowledge base of biologic pathways and processes. Genome Biol. 8, R39 (2007).

Kanehisa, M., Goto, S., Kawashima, S., Okuno, Y. & Hattori, M. The KEGG resource for deciphering the genome. Nucleic Acids Res. 32, D277–80 (2004).

Barrett, T., Wilhite, S. E., Ledoux, P., Evangelista, C., Kim, I. F., Tomashevsky, M., Marshall, K. A., Phillippy, K. H., Sherman, P. M., Holko, M., Yefanov, A., Lee, H., Zhang, N., Robertson, C. L., Serova, N., Davis, S. & Soboleva, A. NCBI GEO: Archive for functional genomics data sets - Update. Nucleic Acids Res. 41, (2013).

Hofree, M., Shen, J. P., Carter, H., Gross, A. & Ideker, T. Network-based stratification of tumor mutations. Nat. Methods 10, 1108–15 (2013).

Segal, E., Shapira, M., Regev, A., Pe’er, D., Botstein, D., Koller, D. & Friedman, N. Module networks: identifying regulatory modules and their condition-specific regulators from gene expression data. Nat. Genet. 34, 166–176 (2003).

Ciriello, G., Miller, M. L., Aksoy, B. A., Senbabaoglu, Y., Schultz, N. & Sander, C. Emerging landscape of oncogenic signatures across human cancers. Nat. Genet. 45, 1127–1133 (2013).

Efron, B., Tibshirani, R., Storey, J. D. & Tusher, V. Empirical Bayes Analysis of a Microarray Experiment. J. Am. Stat. Assoc. 96, 1151–1160 (2001).

Way, G. P., Rudd, J., Wang, C., Hamidi, H., Fridley, B. L., Konecny, G., Goode, E. L., Greene, C. S. & Doherty, J. A. High-grade serous ovarian cancer subtypes are similar across populations. Biorxiv (http://dx.doi.org/10.1101/030239)

Wels, J., Kaplan, R. N., Rafii, S. & Lyden, D. Migratory neighbors and distant invaders: tumor-associated niche cells. Genes Dev. 22, 559–74 (2008).

Le Blanc, K. & Mougiakakos, D. Multipotent mesenchymal stromal cells and the innate immune system. Nat. Rev. Immunol. 12, 383–396 (2012).

Heuvers, M. E., Aerts, J. G., Cornelissen, R., Groen, H., Hoogsteden, H. C. & Hegmans, J. P. Patient-tailored modulation of the immune system may revolutionize future lung cancer treatment. BMC Cancer 12, 580 (2012).

Liu, H., Ma, Q., Xu, Q., Lei, J., Li, X., Wang, Z. & Wu, E. Therapeutic potential of perineural invasion, hypoxia and desmoplasia in pancreatic cancer. Curr. Pharm. Des. 18, 2395–403 (2012).

Riester, M., Wei, W., Waldron, L., Culhane, A. C., Trippa, L., Oliva, E., Kim, S.-H., Michor, F., Huttenhower, C., Parmigiani, G. & Birrer, M. J. Risk prediction for late-stage ovarian cancer by metaanalysis of 1525 patient samples. J. Natl. Cancer Inst. 106, dju048-(2014).

Tothill, R. W., Tinker, A. V., George, J., Brown, R., Fox, S. B., Lade, S., Johnson, D. S., Trivett, M. K., Etemadmoghadam, D., Locandro, B., Traficante, N., Fereday, S., Hung, J. A., Chiew, Y. E., Haviv, I., Gertig, D., Defazio, A. & Bowtell, D. D. L. Novel molecular subtypes of serous and endometrioid ovarian cancer linked to clinical outcome. Clin. Cancer Res. 14, 5198–5208 (2008).

Puisieux, A., Brabletz, T. & Caramel, J. Oncogenic roles of EMT-inducing transcription factors. Nat. Cell Biol. 16, 488–94 (2014).

Ilic, D., Furuta, Y., Kanazawa, S., Takeda, N., Sobue, K., Nakatsuji, N., Nomura, S., Fujimoto, J., Okada, M. & Yamamoto, T. Reduced cell motility and enhanced focal adhesion contact formation in cells from FAK-deficient mice. Nature 377, 539–44 (1995).

Chautard, E., Fatoux-Ardore, M., Ballut, L., Thierry-Mieg, N. & Ricard-Blum, S. MatrixDB, the extracellular matrix interaction database. Nucleic Acids Res. 39, D235–40 (2011).

Barker, T. H., Baneyx, G., Cardó-Vila, M., Workman, G. A., Weaver, M., Menon, P. M., Dedhar, S., Rempel, S. A., Arap, W., Pasqualini, R., Vogel, V. & Sage, E. H. SPARC regulates extracellular matrix organization through its modulation of integrin-linked kinase activity. J. Biol. Chem. 280, 36483–93 (2005).

Dong, J., Grunstein, J., Tejada, M., Peale, F., Frantz, G., Liang, W.-C., Bai, W., Yu, L., Kowalski, J.,Liang, X., Fuh, G., Gerber, H.-P. & Ferrara, N. VEGF-null cells require PDGFR alpha signaling-mediated stromal fibroblast recruitment for tumorigenesis. EMBO J. 23, 2800–10 (2004).

Singh, A. & Settleman, J. EMT, cancer stem cells and drug resistance: an emerging axis of evil in the war on cancer. Oncogene 29, 4741–51 (2010).

Seton-Rogers, S. Layers of regulation. Nat. Rev. Cancer 11, 689 (2011).

Yang, J. & Weinberg, R. A. Epithelial-mesenchymal transition: at the crossroads of development and tumor metastasis. Dev. Cell 14, 818–29 (2008).

Gentles, A. J., Plevritis, S. K., Majeti, R. & Alizadeh, A. A. Association of a leukemic stem cell gene expression signature with clinical outcomes in acute myeloid leukemia. JAMA 304, 2706–15 (2010).

Lopez-Novoa, J. M. & Nieto, M. A. Inflammation and EMT: an alliance towards organ fibrosis and cancer progression. EMBO Mol. Med. 1, 303–14 (2009).

Steinmetz, R., Wagoner, H. A., Zeng, P., Hammond, J. R., Hannon, T. S., Meyers, J. L. & Pescovitz, O. H. Mechanisms regulating the constitutive activation of the extracellular signal-regulated kinase (ERK) signaling pathway in ovarian cancer and the effect of ribonucleic acid interference for ERK1/2 on cancer cell proliferation. Mol. Endocrinol. 18, 2570–82 (2004).

Tamura, R. E., de Vasconcellos, J. F., Sarkar, D., Libermann, T. A., Fisher, P. B. & Zerbini, L. F. GADD45 proteins: central players in tumorigenesis. Curr. Mol. Med. 12, 634–51 (2012).

Tetreault, M.-P., Yang, Y. & Katz, J. P. Krüppel-like factors in cancer. Nat. Rev. Cancer 13, 701–13 (2013).

Gentles, A. J., Newman, A. M., Liu, C. L., Bratman, S. V, Feng, W., Kim, D., Nair, V. S., Xu, Y., Khuong, A., Hoang, C. D., Diehn, M., West, R. B., Plevritis, S. K. & Alizadeh, A. a. The prognostic landscape of genes and infiltrating immune cells across human cancers. Nat. Med. (2015). doi:10.1038/nm.3909

The Cancer Genome Atlas Research Network. Comprehensive genomic characterization defines human glioblastoma genes and core pathways. Nature 455, 1061–1068 (2008).

Gravendeel, L. A. M., Kouwenhoven, M. C. M., Gevaert, O., de Rooi, J. J., Stubbs, A. P., Duijm, J. E., Daemen, A., Bleeker, F. E., Bralten, L. B. C., Kloosterhof, N. K., De Moor, B., Eilers, P. H. C., Van der Spek, P. J., Kros, J. M., Sillevis Smitt, P. A. E., van den Bent, M. J. & French, P. J. Intrinsic gene expression profiles of gliomas are a better predictor of survival than histology. Cancer Res. 69, 9065–72 (2009).

Collisson, E. a., Campbell, J. D., Brooks, A. N., Berger, A. H., Lee, W., Chmielecki, J., Beer, D. G., Cope, L., Creighton, C. J., Danilova, L., Ding, L., Getz, G., Hammerman, P. S., Neil Hayes, D., Hernandez, B., Herman, J. G., Heymach, J. V., Jurisica, I., Kucherlapati, R., Kwiatkowski, D., Ladanyi, M., Robertson, G., Schultz, N., Shen, R., Sinha, R., Sougnez, C., Tsao, M.-S., Travis, W. D., Weinstein, J. N., Wigle, D. a., Wilkerson, M. D., Chu, A., Cherniack, A. D., Hadjipanayis, A., Rosenberg, M., Weisenberger, D. J., Laird, P. W., Radenbaugh, A., Ma, S., Stuart, J. M., Averett Byers, L., Baylin, S. B., Govindan, R., Meyerson, M., Gabriel, S. B., Cibulskis, K., Kim, J., Stewart, C., Lichtenstein, L., Lander, E. S., Lawrence, M. S., Kandoth, C., Fulton, R., Fulton, L. L., McLellan, M. D., Wilson, R. K., Ye, K., Fronick, C. C., Maher, C. a., Miller, C. a., Wendl, M. C., Cabanski, C., Mardis, E., Wheeler, D., Balasundaram, M., Butterfield, Y. S. N., Carlsen, R., Chuah, E., Dhalla, N., Guin, R., Hirst, C., Lee, D., Li, H. I., Mayo, M., Moore, R. a., Mungall, A. J., Schein, J. E., Sipahimalani, P., Tam, A., Varhol, R., Gordon Robertson, a., Wye, N., Thiessen, N., Holt, R. a., Jones, S. J. M., Marra, M. a., Imielinski, M., Onofrio, R. C., Hodis, E., Zack, T., Helman, E., Sekhar Pedamallu, C., Mesirov, J., Saksena, G., Schumacher, S. E., Carter, S. L., Garraway, L., Beroukhim, R., Lee, S., Mahadeshwar, H. S., Pantazi, A., Protopopov, A., Ren, X., Seth, S., Song, X., Tang, J., Yang, L., Zhang, J., Chen, P.-C., Parfenov, M., Wei Xu, A., Santoso, N., Chin, L., Park, P. J., Hoadley, K. a., Todd Auman, J., Meng, S., Shi, Y., Buda, E., Waring, S., Veluvolu, U., Tan, D., Mieczkowski, P. a., Jones, C. D., Simons, J. V., Soloway, M. G., Bodenheimer, T., Jefferys, S. R., Roach, J., Hoyle, A. P., Wu, J., Balu, S., Singh, D., Prins, J. F., Marron, J. S., Parker, J. S., Perou, C. M., Liu, J., Maglinte, D. T., Lai, P. H., Bootwalla, M. S., van Den Berg, D. J., Triche Jr, T., Cho, J., DiCara, D., Heiman, D., Lin, P., Mallard, W., Voet, D., Zhang, H., Zou, L., Noble, M. S., Gehlenborg, N., Thorvaldsdottir, H., Nazaire, M.-D., Robinson, J., Arman Aksoy, B., Ciriello, G., Taylor, B. S., Dresdner, G., Gao, J., Gross, B., Seshan, V. E., Reva, B., Onur Sumer, S., Weinhold, N., Sander, C., Ng, S., Zhu, J., Benz, C. C., Yau, C., Haussler, D., Spellman, P. T., Kimes, P. K., Broom, B. M., Wang, J., Lu, Y., Kwok Shing Ng, P., Diao, L., Liu, W., Amos, C. I., Akbani, R., Mills, G. B., Curley, E., Paulauskis, J., Lau, K., Morris, S., Shelton, T., Mallery, D., Gardner, J., Penny, R., Saller, C., Tarvin, K., Richards, W. G., Cerfolio, R., Bryant, A., Daniel P. Raymond,:, Pennell, N. a., Farver, C., Czerwinski, C., Huelsenbeck-Dill, L., Iacocca, M., Petrelli, N., Rabeno, B., Brown, J., Bauer, T., Dolzhanskiy, O., Potapova, O., Rotin, D., Voronina, O., Nemirovich-Danchenko, E., Fedosenko, K. V., Gal, A., Behera, M., Ramalingam, S. S., Sica, G., Flieder, D., Boyd, J., Weaver, J., Kohl, B., Huy Quoc Thinh, D., Sandusky, G., Juhl, H., Duhig, E., Illei, P., Gabrielson, E., Shin, J., Lee, B., Rogers, K., Trusty, D., Brock, M. V., Williamson, C., Burks, E., Rieger-Christ, K., Holway, A., Sullivan, T., Asiedu, M. K., Kosari, F., Rekhtman, N., Zakowski, M., Rusch, V. W., Zippile, P., Suh, J., Pass, H., Goparaju, C., Owusu-Sarpong, Y., Bartlett, J. M. S., Kodeeswaran, S., Parfitt, J., Sekhon, H., Albert, M., Eckman, J., Myers, J. B., Cheney, R., Morrison, C., Gaudioso, C., Borgia, J. a., Bonomi, P., Pool, M., Liptay, M. J., Moiseenko, F., Zaytseva, I., Dienemann, H., Meister, M., Schnabel, P. a., Muley, T. R., Peifer, M., Gomez-Fernandez, C., Herbert, L., Egea, S., Huang, M., Thorne, L. B., Boice, L., Hill Salazar, A., Funkhouser, W. K., Kimryn Rathmell, W., Dhir, R., Yousem, S. a., Dacic, S., Schneider, F., Siegfried, J. M., Hajek, R., Watson, M. a., McDonald, S., Meyers, B., Clarke, B., Yang, I. a., Fong, K. M., Hunter, L., Windsor, M., Bowman, R. V., Peters, S., Letovanec, I., Khan, K. Z., Jensen, M. a., Snyder, E. E., Srinivasan, D., Kahn, A. B.,Baboud, J., Pot, D. a., Mills Shaw, K. R., Sheth, M., Davidsen, T., Demchok, J. a., Yang, L., Wang, Z., Tarnuzzer, R., Claude Zenklusen, J., Ozenberger, B. a. & Sofia, H. J. Comprehensive molecular profiling of lung adenocarcinoma. Nature 511, 543–50 (2014).

Gentles, A. J., Alizadeh,A. A., Lee, S. I., Myklebust, J. H., Shachaf, C. M., Shahbaba, B., Levy, R., Koller, D. & Plevritis, S. K. A pluripotency signature predicts histologic transformation and influences survival in follicular lymphoma patients. Blood 114, 3158–3166 (2009).

Chen, F., Kook, H., Milewski, R., Gitler, A. D., Lu, M. M., Li, J., Nazarian, R., Schnepp, R., Jen, K., Biben, C.,Runke, G., Mackay, J. P., Novotny, J., Schwartz, R. J., Harvey, R. P., Mullins, M. C. & Epstein, J. A. Hop is an unusual homeobox gene that modulates cardiac development. Cell 110, 713–723 (2002).

Kee, H. J., Kim, J.-R., Nam, K.-I., Park, H. Y., Shin, S., Kim, J. C., Shimono, Y., Takahashi, M., Jeong, M. H., Kim, N., Kim, K. K. & Kook, H. Enhancer of polycomb1, a novel homeodomain only protein-binding partner, induces skeletal muscle differentiation. J. Biol. Chem. 282, 7700–9 (2007).

Kook, H., Lepore, J. J., Gitler, A. D., Lu, M. M., Yung, W. W. M., Mackay, J., Zhou, R., Ferrari, V., Gruber, P. & Epstein, J. a. Cardiac hypertrophy and histone deacetylase-dependent transcriptional repression mediated by the atypical homeodomain protein Hop. J. Clin. Invest. 112, 863–871 (2003).

Katoh, H., Yamashita, K., Waraya, M., Margalit, O., Ooki, A., Tamaki, H., Sakagami, H., Kokubo, K., Sidransky, D. & Watanabe, M. Epigenetic Silencing of HOPX Promotes Cancer Progression in Colorectal Cancer. Neoplasia 14, 559–IN6 (2012).

Waraya, M., Yamashita,K., Katoh, H., Ooki, A., Kawamata, H., Nishimiya, H., Nakamura, K., Ema, A. & Watanabe, M. Cancer specific promoter CpG Islands hypermethylation of HOP homeobox (HOPX) gene and its potential tumor suppressive role in pancreatic carcinogenesis. BMC Cancer 12, 397 (2012).

Chen, Y., Yang, L., Cui,T., Pacyna-Gengelbach, M. & Petersen, I. HOPX is methylated and exerts tumour suppressive function through Ras-induced senescence in human lung cancer. J. Pathol. (2014). doi:10.1002/path.4469

Jain, R., Li, D., Gupta, M., Manderfield, L. J., Ifkovits, J. L., Wang, Q., Liu, F., Liu, Y., Poleshko, A., Padmanabhan, A., Raum, J. C., Li, L., Morrisey, E. E., Lu, M. M., Won, K.-J. & Epstein, J. A. Integration of Bmp and Wnt signaling by Hopx specifies commitment of cardiomyoblasts. Science (80-.). 348, aaa6071–aaa6071 (2015).

Logsdon, B. a., Gentles,a. J., Miller, C. P., Blau, C. a., Becker, P. S. & Lee, S.-I. Sparse expression bases in cancer reveal tumor drivers. Nucleic Acids Res. 1–13 (2015). doi:10.1093/nar/gku1290

Kim, J.-A., Choi, H.-K., Kim, T.-M., Leem, S.-H. & Oh, I.-H. Regulation of mesenchymal stromal cells through fine tuning of canonical Wnt signaling. Stem Cell Res. 14, 356–368 (2015).

Macheda, M. L. & Stacker, S. a. Importance of Wnt signaling in the tumor stroma microenvironment. Curr. Cancer Drug Targets 8, 454–65 (2008).

Savagner, P., Yamada, K. M. & Thiery, J. P. The zinc-finger protein slug causes desmosome dissociation, an initial and necessary step for growth factor-induced epithelial-mesenchymal transition. J. Cell Biol. 137, 1403–19 (1997).

Mishra, P. J., Mishra, P. J., Humeniuk, R., Medina, D. J., Alexe, G., Mesirov, J. P., Ganesan, S., Glod, J. W. &Banerjee, D. Carcinoma-associated fibroblast-like differentiation of human mesenchymal stem cells. Cancer Res. 68, 4331–4339 (2008).

Li, N., Yousefi, M., Nakauka-Ddamba, A., Jain, R., Tobias, J., Epstein, J. A., Jensen, S. T. & Lengner, C. J. Single-Cell Analysis of Proxy Reporter Allele-Marked Epithelial Cells Establishes Intestinal Stem Cell Hierarchy. Stem Cell Reports 3, 876–91 (2014).

Jain, R., Barkauskas, C. E., Takeda, N., Bowie, E. J., Aghajanian, H., Wang, Q., Padmanabhan, A., Manderfield, L. J., Gupta, M., Li, D., Li, L., Trivedi, C. M., Hogan, B. L. M. & Epstein, J. A. Plasticity of Hopx(+) type I alveolar cells to regenerate type II cells in the lung. Nat. Commun. 6, 6727 (2015).

Ma, S., Xie, N., Li, W., Yuan, B., Shi, Y. & Wang, Y. Immunobiology of mesenchymal stem cells. Cell Death Differ. 21, 216–25 (2014).

Karnoub, A. E., Dash, A. B., Vo, A. P., Sullivan, A., Brooks, M. W., Bell, G. W., Richardson, A. L., Polyak, K., Tubo, R. & Weinberg, R. a. Mesenchymal stem cells within tumour stroma promote breast cancer metastasis. Nature 449, 557–563 (2007).

Law, C. W., Chen, Y., Shi, W. & Smyth, G. K. voom: Precision weights unlock linear model analysis tools for RNA-seq read counts. Genome Biol. 15, R29 (2014).

http://gdac.broadinstitute.org/runs/stddata2014_03_16/data/OV/20140316/.

Johnson, W. E., Li, C. & Rabinovic, A. Adjusting batch effects in microarray expression data using empirical Bayes methods. Biostatistics 8, 118–127 (2007).

Denkert, C., Budczies, J., Darb-Esfahani, S., Gyorffy, B., Sehouli, J., Könsgen, D., Zeillinger, R., Weichert, W., Noske, A., Buckendahl, A. C., Müller, B. M., Dietel, M. & Lage, H. A prognostic gene expression index in ovarian cancer - Validation across different independent data sets. J. Pathol. 218, 273–280 (2009).

Bonome, T., Levine, D. A., Shih, J., Randonovich, M., Pise-Masison, C. A., Bogomolniy, F., Ozbun, L., Brady, J., Barrett, J. C., Boyd, J. & Birrer, M. J. A gene signature predicting for survival in suboptimally debulked patients with ovarian cancer. Cancer Res. 68, 5478–5486 (2008).

Hendrix, N. D., Wu, R., Kuick, R., Schwartz, D. R., Fearon, E. R. & Cho, K. R. Fibroblast growth factor 9 has oncogenic activity and is a downstream target of Wnt signaling in ovarian endometrioid adenocarcinomas. Cancer Res. 66, 1354–1362 (2006).

Mok, S. C., Bonome, T., Vathipadiekal, V., Bell, A., Johnson, M. E., Wong, kwong kwok, Park, D. C., Hao, K., Yip, D. K. P., Donninger, H., Ozbun, L., Samimi, G., Brady, J., Randonovich, M., Pise-Masison, C.A., Barrett, J. C., Wong, W. H., Welch, W. R., Berkowitz, R. S. & Birrer, M. J. A Gene Signature Predictive for Outcome in Advanced Ovarian Cancer Identifies a Survival Factor: Microfibril-Associated Glycoprotein 2. Cancer Cell 16, 521–532 (2009).

Meyniel, J.-P., Cottu, P. H., Decraene, C., Stern, M.-H., Couturier, J., Lebigot, I., Nicolas, A., Weber, N., Fourchotte, V., Alran, S., Rapinat, A., Gentien, D., Roman-Roman, S., Mignot, L. & Sastre-Garau, X. A genomic and transcriptomic approach for a differential diagnosis between primary and secondary ovarian carcinomas in patients with a previous history of breast cancer. BMC Cancer 10, 222 (2010).

Ferriss, J. S., Kim, Y., Duska, L., Birrer, M., Levine, D. A., Moskaluk, C., Theodorescu, D. & Lee, J. K. Multi-gene expression predictors of single drug responses to adjuvant chemotherapy in ovarian carcinoma: Predicting platinum resistance. PLoS One 7, (2012).

Gautier, L., Cope, L., Bolstad, B. M. & Irizarry, R. A. Affy – Analysis of Affymetrix GeneChip data at the probe level. Bioinformatics 20, 307–315 (2004).

Maglott, D., Ostell, J., Pruitt, K. D. & Tatusova, T. Entrez gene: Gene-centered information at NCBI. Nucleic Acids Res. 39, (2011).

Dai, M., Wang, P., Boyd, A. D., Kostov, G., Athey, B., Jones, E. G., Bunney, W. E., Myers, R. M., Speed, T. P., Akil, H., Watson, S. J. & Meng, F. Evolving gene/transcript definitions significantly alter the interpretation of GeneChip data. Nucleic Acids Res. 33, (2005).

Tibshirani, R. Regression Selection and Shrinkage via the Lasso. J. R. Stat. Soc. B 58, 267–288 (1994).

Tibshirani, R., Bien, J., Friedman, J., Hastie, T., Simon, N., Taylor, J. & Tibshirani, R. J. Strong rules for discarding predictors in lasso-type problems. J. R. Stat. Soc. Ser. B Stat. Methodol. 74, 245–266 (2012).

Marquardt, D. W. & Snee, R. D. Ridge Regression in Practice. Source Am. Stat. 29, 3–20 (1975).

Sardy, S. On the practice of rescaling covariates. Int. Stat. Rev. 76, 285–297 (2008).

de los Campos, G., Hickey, J. M., Pong-Wong, R., Daetwyler, H. D. & Calus, M. P. L.Whole-genome regression and prediction methods applied to plant and animal breeding. Genetics 193,327–345 (2013).

Schmidt, M., Niculescu-Mizil, A. & Murphy, K. P. Learning Graphical Model Structure using L1-Regularization Paths. Proc. AAAI Conf. Artif. Intell. 22, 1278 (2007).

Mu, B. & How, J. P. Learning Sparse Gaussian Graphical Model with L 0-regularization. Tech. Rep. 1–13 (2014).

Friedman, J., Hastie, T. & Tibshirani, R. Applications of the lasso and grouped lasso to the estimation of sparse graphical models. Tech. Rep. 1–22 (2010).

Lee, S.-I., Pe’er, D., Dudley, A. M., Church, G. M. & Koller, D. Identifying regulatory mechanisms using individual variation reveals key role for chromatin modification. Proc. Natl. Acad. Sci. U. S. A. 103, 14062–14067 (2006).

Lee, S. I., Dudley, A. M., Drubin, D., Silver, P. A., Krogan, N.J., Pe’er, D. & Koller, D. Learning a prior on regulatory potential from eQTL data. PLoS Genet. 5, (2009).

Akavia, U. D., Litvin, O., Kim, J., Sanchez-Garcia, F., Kotliar,D., Causton, H. C., Pochanard, P., Mozes, E., Garraway, L. A. & Pe’Er, D. An integrated approach to uncover drivers of cancer. Cell 143, 1005–1017 (2010).

cBio Cancer Genomics Portal: http://cbioportal.org.

Mermel, C. H., Schumacher, S. E., Hill, B., Meyerson, M. L., Beroukhim, R. & Getz, G. GISTIC2.0 facilitates sensitive and confident localization of the targets of focal somatic copy-number alteration in human cancers. Genome Biol. 12, R41 (2011).

Mardia, K. V, Kent, J. T. & Bibby, J. M. Multivariate Analysis. (Academic Press, 1979). at <http://www.amazon.com/dp/0124712525>

Lauritzen, S. L. Graphical Models. (Oxford University Press, 1996).

Tseng, P. Convergence of a block coordinate descent method for nondifferentiable minimization. J. Optim. Theory Appl. 109, 475–494 (2001).

Josse, J., Chavent, M., Liquet, B. & Husson, F. Handling Missing Values with Regularized Iterative Multiple Correspondence Analysis. J. Classif. 29, 91–116 (2012).

Witten, D. M., Friedman, J. H. & Simon, N. New Insights and Faster Computations for the Graphical Lasso. J. Comput. Graph. Stat. 20, 892–900 (2011).

Tibshirani, R. Regression shrinkage and selection via the lasso. J. R. Stat. Soc. Ser. B267–288 (1996).

ImageJ: http://imagej.nih.gov/ij/.

Gene Expression Omnibus (GEO): http://www.ncbi.nlm.nih.gov/geo/.

